# Discovery of Semicarbazone and Thiosemicarbazone Analogs as Competitive SARS-CoV-2 Virus Main Protease (M^pro^) Inhibitors

**DOI:** 10.64898/2026.03.11.711159

**Authors:** Luiza Valença Barreto, Estela Mariana Guimarães Lourenço, Elany Barbosa da Silva, Mariana O. de Godoy, Luan Carvalho Martins, Mariana Laureano de Souza, Renata G. Almeida, Vitor Leonardo Silva Cunha, Magno Carvalho Pires, Stefânia Neiva Lavorato, Thiago Belarmino de Souza, Ana Carolina Oliveira Bretas, Flaviano Melo Ottoni, Eufrânio N. da Silva Júnior, Glaucius Oliva, Ricardo José Alves, Renata Barbosa De Oliveira, Rafael Victorio Carvalho Guido, Rafaela Salgado Ferreira

## Abstract

Despite the development of vaccines and antivirals, coronavirus disease 2019 (COVID-19) continues to affect populations worldwide. Given the high mutation rate of the SARS-CoV-2 virus and reports of drug resistance, there is a continued need for new therapeutic options. SARS-CoV-2 main protease (M^pro^) is essential for viral replication and is a conserved target among coronaviruses. Most known M^pro^ inhibitors target the active site, although allosteric sites have already been identified. In this study, we conducted a virtual screening of 2,060 compounds targeting an allosteric site of SARS-CoV-2 M^pro^. From this screen, 41 computational hits and analogs were selected and evaluated using biochemical assays against SARS-CoV-2 M^pro^. Among them, compound **25**, a semicarbazone, demonstrated a half-maximal inhibitory concentration (IC_50_) of 99 μM. Additionally, two thiosemicarbazone analogs (compounds **50** and **51**) inhibited SARS-CoV-2 M^pro^ with IC_50_ values of 61 μM and 70 μM. Biochemical assays suggest that these compounds act as noncovalent competitive inhibitors of SARS-CoV-2 M^pro^. Molecular dynamics simulations revealed that compound **25** is unstable at the allosteric site of SARS-CoV-2 M^pro^ but forms stable and favorable interactions at the active site, supporting its potential as a competitive inhibitor, a finding subsequently confirmed by biochemical assays. Our structure-based computational and biochemical approach identified semicarbazone and thiosemicarbazone scaffolds as promising candidates for the development of reversible SARS-CoV-2 M^pro^ inhibitors.

## 1. Introduction

The first cases of COVID-19 were reported in late 2019 in Wuhan, China. In 2020, the World Health Organization (WHO) declared the outbreak a pandemic, marking an unprecedented crisis in global public health.^1–3^ COVID-19 is caused by a highly infectious β-coronavirus, the severe acute respiratory syndrome coronavirus 2 (SARS-CoV-2).^3–5^ By August 2025, approximately 778 million cases and over 7 million deaths had been reported worldwide.^6^ Over the past five years, substantial progress has been made in managing the disease, primarily through the development of vaccines and antiviral therapies.^7,8^ Nonetheless, there remains an urgent need for safe, effective, and affordable therapeutic options.^8–10^

The SARS-CoV-2 genome exhibits high similarity to other coronaviruses, sharing 79% sequence identity with SARS-CoV, the causative agent of severe acute respiratory syndrome, and approximately 50% with MERS-CoV, which causes Middle East respiratory syndrome (MERS).^11–14^ SARS-CoV-2 possesses a positive-sense, single-stranded RNA genome of approximately 30 kb, which encodes open reading frames (ORFs) ORF1a and ORF1b. Their translation results in two overlapping polyproteins, pp1a and pp1ab.^15,16^ These polyproteins are cleaved by the SARS-CoV-2 Main Protease (M^pro^, also known as 3-chymotrypsin-like protease, 3CLpro), yielding 11 non-structural viral proteins essential for viral replication.^17–19^ SARS-CoV-2 M^pro^ is a 33.8-kDa cysteine protease that shares 96% protein sequence identity with SARS-CoV M^pro^, 50.7% with MERS-CoV M^pro^, and 99% with M^pro^ from bat coronaviruses.^20,21^ This high degree of conservation across several coronaviruses makes M^pro^ an attractive target for the development of pan-coronavirus inhibitors.^22,23^

Structurally, SARS-CoV-2 M^pro^ is a homodimer, with each monomer comprising three domains that adopt a chymotrypsin fold similar to 3C-like proteases from the picornavirus family.^24^ The enzymés active site comprises a catalytic dyad, Cys145 and His41, positioned within a shallow cleft between domains I and II. Domain III is α-helical and plays a critical role in dimerization and enzymatic activity. Substrate recognition occurs specifically with glutamine residues in P1, hydrophobic residues in P2, and serine and alanine in P1’. Notably, no human protease shares this substrate cleavage specificity and sequence identity, enhancing the potential for developing selective and safe drugs.^17,23,25^ Given the essential role of SARS-CoV-2 M^pro^ in viral replication and its high sequence conservation, several covalent, non-covalent small-molecules, as well as peptide-based inhibitors, that bind to its active site have been identified and have demonstrated antiviral efficacy in cell culture and animal models.^25–30^

Structure-based drug design (SBDD) strategies have played a key role in the development of these drugs, as exemplified by the development of nirmatrelvir (administered in combination with ritonavir as Paxlovid™), a reversible covalent inhibitor of SARS-CoV-2 M^pro^ (IC_50_ = 0.0192 µM)^31–36^, and ensitrelvir, a non-peptide and non-covalent inhibitor (IC_50_ = 0.013 μM) optimized from a virtual screening hit (Figure 1).^37,38^ A second generation of SARS-CoV-2 M^pro^ inhibitors comprises several approved medicines and compounds currently undergoing clinical trials. Examples include nirmatrelvir analogs such as PF-07817883 (ibuzatrelvir), a covalent inhibitor developed by Pfizer that eliminates the need for co-administration with ritonavir.^39,40^ Additional examples of covalent SARS-CoV-2 M^pro^ inhibitors approved for the treatment of patients with mild to moderate COVID-19 are atilotrevir (GST-HG171)^41^ and simnotrelvir (SIM-0417)^42,43^, both co-administered with ritonavir, and leritrelvir (RAY1216), which does not require co-administration with ritonavir (Figure 1).^44,45^

**Figure 1.**
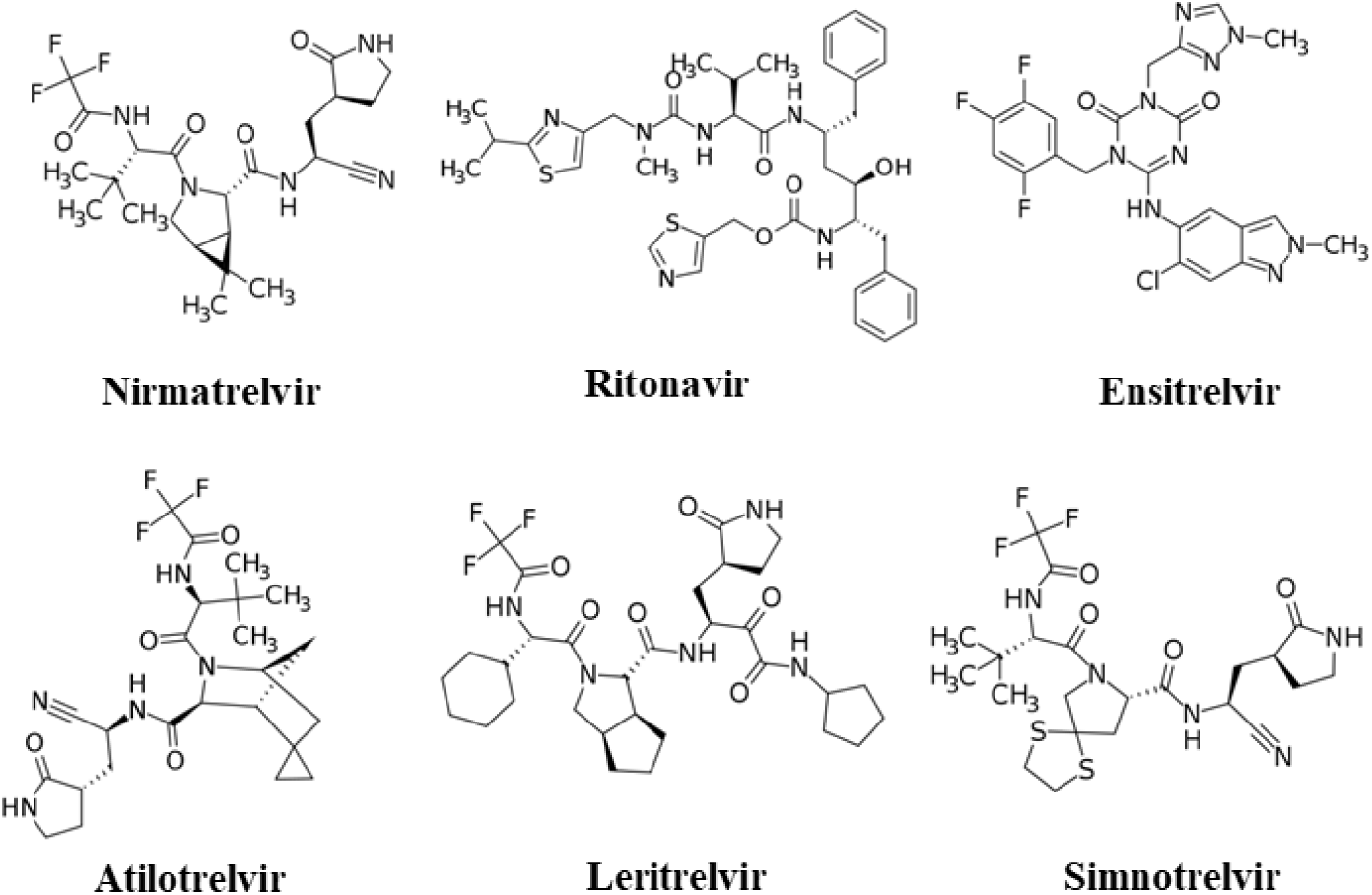
Inhibitors that target SARS-CoV-2 M^pro^ have been approved for use. 2D chemical structures generated in Marvin Sketch.

Despite the available treatments for COVID-19, the continued emergence of SARS-CoV-2 variants raises significant concerns about resistance to current antiviral therapies.^10,19,46^ Mutations within the SARS-CoV-2 M^pro^ active site that directly affect the inhibition by nirmatrelvir have been reported.^47–49^ Naturally occurring mutations at residues Ser144, Met165, Glu166, His172, and Gln192 have been shown to alter viral replication kinetics and confer drug resistance. Among these, Glu166 has been recognized as a critical hotspot; notably, the Leu50Phe/Glu166Val double mutant resulted in an 80-fold increase in resistance to nirmatrelvir in antiviral assays.^48^ Additional mutations - Ser144Leu, Met165Tyr, His172Tyr, and Gln192Thr – not only disrupt key interactions between nirmatrelvir and SARS-CoV-2 M^pro^ but also impact enzymatic function, leading to up to a 127-fold increase in the inhibition constant (*K_i_*).^47^ Resistance-associated mutations have also been observed at positions outside the nirmatrelvir binding pocket, including Gly15Ser, Thr21Ile, Leu89Phe, Lys90Arg, Pro108Ser, Pro132His, and Leu205Val. These substitutions are thought to restore the catalytic efficiency of SARS-CoV-2 M^pro^, thereby contributing indirectly to resistance.^46^ These findings highlight the urgent need to develop additional antiviral agents that remain effective against current and emerging SARS-CoV-2 variants.

One strategy to circumvent resistance to current SARS-CoV-2 M^pro^ inhibitors is to design compounds that target allosteric sites.^50–57^ Gunter *et al.* identified two allosteric sites in SARS-CoV-2 M^pro^. Allosteric site 1 is a hydrophobic pocket located in the C-terminal dimerization domain.^52^ Several compounds have been shown to bind to this site and exhibit antiviral activity. For instance, the anticancer agent pelitinib (PDB code: 7AXM) and the chemokine receptor antagonist RS-102895 (PDB code: 7ABU) demonstrated antiviral activity in Vero E6 cells, with half-maximal effective concentration (EC_50_) = 1.25 µM and 20 µM, respectively. Ifenprodil (PDB code: 7AQI), a vasodilator that has progressed to phase III clinical trials for COVID-19 treatment in hospitalized patients, also binds to SARS-CoV-2 M^pro^ allosteric site 1. Additionally, the anticoagulant apixaban was predicted via molecular docking to bind this site and showed potent inhibition of SARS-CoV-2 M^pro^ (IC₅₀ = 0.01 µM), along with significant antiviral activity in Calu-3 cells (EC₅₀ = 1.8 µM).^58^ A recent review reported 12 non-competitive SARS-CoV-2 M^pro^ inhibitors that interact with allosteric sites 1, 2, or other alternative pockets.^33^ These findings underscore the therapeutic potential of targeting allosteric sites, particularly those near the dimerization interface, which is essential for SARS-CoV-2 M^pro^ activity.^52^

In the present study, we reported a docking-based virtual screening aiming at identifying novel SARS-CoV-2 M^pro^ inhibitors from a library of 2,060 compounds.^5,59^

## 2. Results and Discussion

### 2.1 Redocking, Cross-Docking, and Virtual Screening against SARS-CoV-2 M^pro^

To select an appropriate structure for molecular docking, we analyzed six SARS-CoV-2 M^pro^ structures with ligands bound to allosteric sites, available in the Protein Data Bank (PDB). Alignment of the five available complexes containing ligands bound to allosteric site 1 (PDBs: 7AQI, 7AMJ, 7ABU, 7APH, and 7AXM) revealed a high degree of structural similarity in the protein conformation surrounding this site (Figure S1). Based on this analysis, we selected the highest-resolution structure (1.70 Å, PDB: 7AQI) for our initial molecular docking experiments at allosteric site 1. For allosteric site 2, only one complex structure was available (1.68 Å, PDB: 7AGA).

We evaluated the performance of two molecular docking software, AutoDock Vina^60^ and the DockThor server^61^, based on redocking and cross-docking experiments (Tables S1 and S2). For allosteric site 1 (Figure 2A), redocking of ifenprodil using DockThor yielded a root-mean-square deviation (RMSD) of 2.8 Å. In cross-docking experiments, RMSD values below 2.0 Å were obtained for PD-168568 and RS-102895 among the top ten scoring poses (Table S1), indicating a reasonable predictive accuracy. However, for pelitinib and tofogliflozin, none of the top ten poses achieved an RMSD ≤ 2.0 Å relative to their crystallographic binding modes, likely due to the solvent-exposed regions of these ligands, which complicate the docking accuracy. During the redocking of ifenprodil, better overlap was observed between the phenolic ring and the crystallographic pose within the hydrophobic pocket of allosteric site 1. However, the solvent-exposed portion of the molecule did not align well, contributing to an overall RMSD > 2.0 Å (Figure 2A). Notably, none of the AutoDock Vina-generated poses achieved an RMSD of less than 2.0 Å. For allosteric site 2 (Figure 2B), redocking of AT-7519 using DockThor was successful, yielding an RMSD of 0.6 Å for the second-ranked pose (Table S1), indicating a high accuracy in reproducing the crystallographic binding conformation.

**Figure 2.**
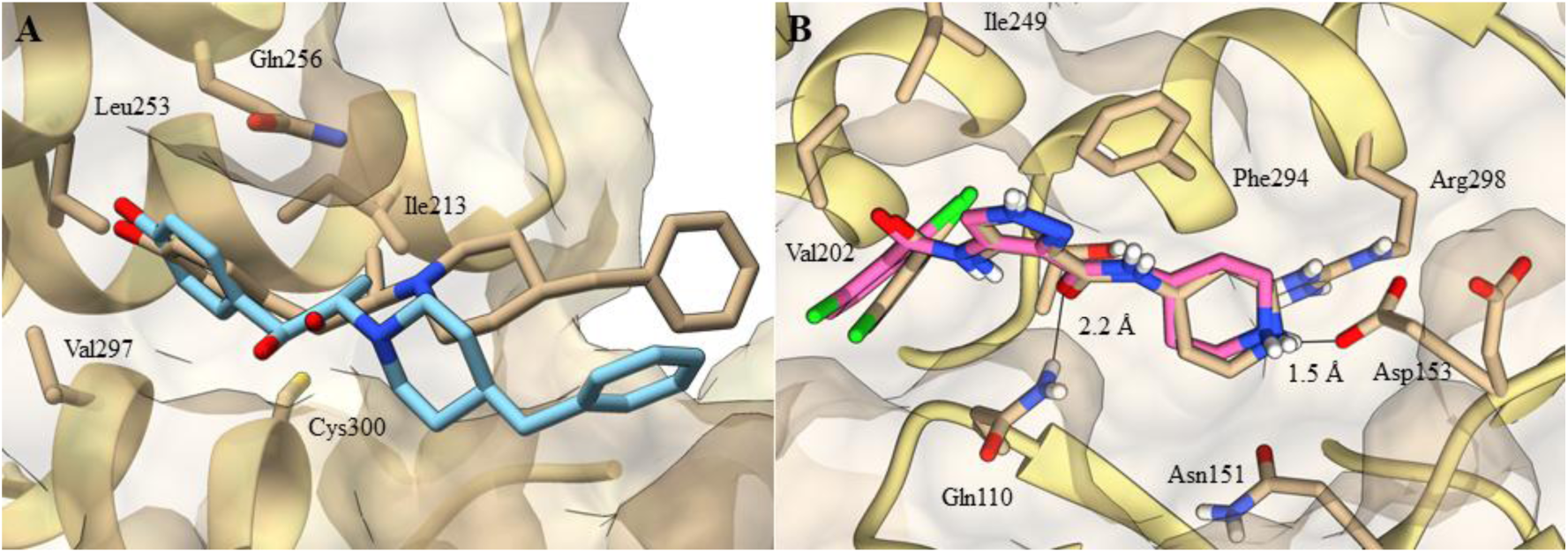
Redocking results at allosteric sites 1 and 2 obtained using the DockThor Server. (A) Redocked pose of ifenprodil (carbons in cyan; RMSD = 2.8 Å) superimposed on its crystallographic binding mode (carbons in beige) at allosteric site 1. (B) Redocked pose of AT-7519 (carbons in pink; RMSD = 0.6 Å) superimposed on its crystallographic binding mode (carbons in beige) at allosteric site 2. The SARS-CoV-2 M^pro^ dimer structures (PDB: 7AQI and 7AGA) are shown as beige surfaces, with key allosteric site residues displayed as sticks. Hydrogen bonds are depicted as black lines, with distances given in Ångströms. Figure generated using UCSF Chimera software.

### 2.2 Discovery of a novel inhibitor through virtual screening

Based on the superior performance observed in redocking and cross-docking experiments, we selected the DockThor server to conduct a virtual screening of two chemical libraries: ENSJ^5^ (884 compounds) and BraCoLi^59^ (1,176 compounds), totaling 2,060 compounds. The docking results were visually inspected, and ligand poses were evaluated based on the overall complementarity to the allosteric binding site and specific interactions with key residues comprising allosteric site 1: Ile213, Leu253, Gln256, Val297, and Cys300.^52^ Compounds with numerous polar groups not involved in favorable polar interactions were excluded. Ultimately, 41 compounds were prioritized for further investigation based on their chemical diversity and favorable interactions within allosteric site 1.

The selected compounds were evaluated in biochemical assays using recombinant SARS-CoV-2 M^pro^, with enzyme activity measured via cleavage of a fluorogenic substrate (Table S3). Two compounds (**21** and **23**) were excluded from testing because of their insolubility in DMSO. Initial inhibition screening was conducted after a 10-minute preincubation of the enzyme with each compound at 100 μM, except for compounds **1**, **2**, **5**, **7**, **14**, **15**, **17**, **20**, **29, 30**, and **33**, which were tested at their maximum soluble concentrations in the assay buffer (25–50 μM). Owing to their intrinsic fluorescence, eight compounds (**4**, **12**, **26**, **34-36**, **38**, and **41**) could not be evaluated by the FRET-based assay. Of the 31 tested compounds, 27 compounds (**1**-**3**, **5**-**11**, **13**, **14**, **17-20**, **22**, **24**, **27-31**, **33**, **37**, **39**, and **40**) showed no significant inhibition of enzyme activity (Table S3). However, four compounds (**15**, **16**, **25,** and **32**) inhibited SARS-CoV-2 M^pro^ activity by at least 50% at 100 µM. Among these, compound **25**, a semicarbazone derivative, exhibited a half-maximal inhibitory concentration (IC_50_) of 99±0.3 μM (Figure S2). The remaining three hits (compounds **15**, **16,** and **32**) exhibited limited solubility at higher concentrations and were not further characterized. The inhibitor nirmatrelvir was used as a positive control and exhibited an IC_50_ of 7.5 ± 0.4 nM (Figure S2).

### 2.3 Molecular dynamics simulations and experimental assays revealed compound 25 as a competitive M^pro^ inhibitor

Following the identification of compound **25** as a SARS-CoV-2 M^pro^ inhibitor, we conducted molecular dynamics (MD) simulations to evaluate the stability of its predicted binding poses within allosteric site 1 and to propose a plausible binding mode. DockThor offers two docking modes: standard mode, which enables exhaustive conformational sampling for detailed analyses, and virtual screening mode, optimized for high-throughput screening of large compound libraries.^62,63^ As the virtual screening mode performs fewer sampling runs than the standard protocol, the best-ranked pose used to select compound **25** may not correspond to the most representative binding conformation. Therefore, we performed molecular dynamics simulations using the highest-ranked poses obtained from both docking strategies implemented in DockThor (Figure S3A and S3B).

Although the standard mode of DockThor employs exhaustive sampling, the top-ranked pose generated by this protocol did not remain stably bound at the M^pro^ allosteric site during the simulations (Supplementary Material, Figure S3C). In contrast, the lowest-energy pose obtained from the VS mode, initially docked to protomer A, underwent a conformational transition and established interactions with the allosteric site of protomer B approximately 50 ns into the simulation. Given this unexpected behavior, we extended the simulation to further explore the interactions of compound **25** with protomer B. Notably, by 200 ns, the compound migrated to the active site of protomer B, where it formed stable interactions and remained bound until the end of the simulation (500 ns) (Figure 3A).

**Figure 3.**
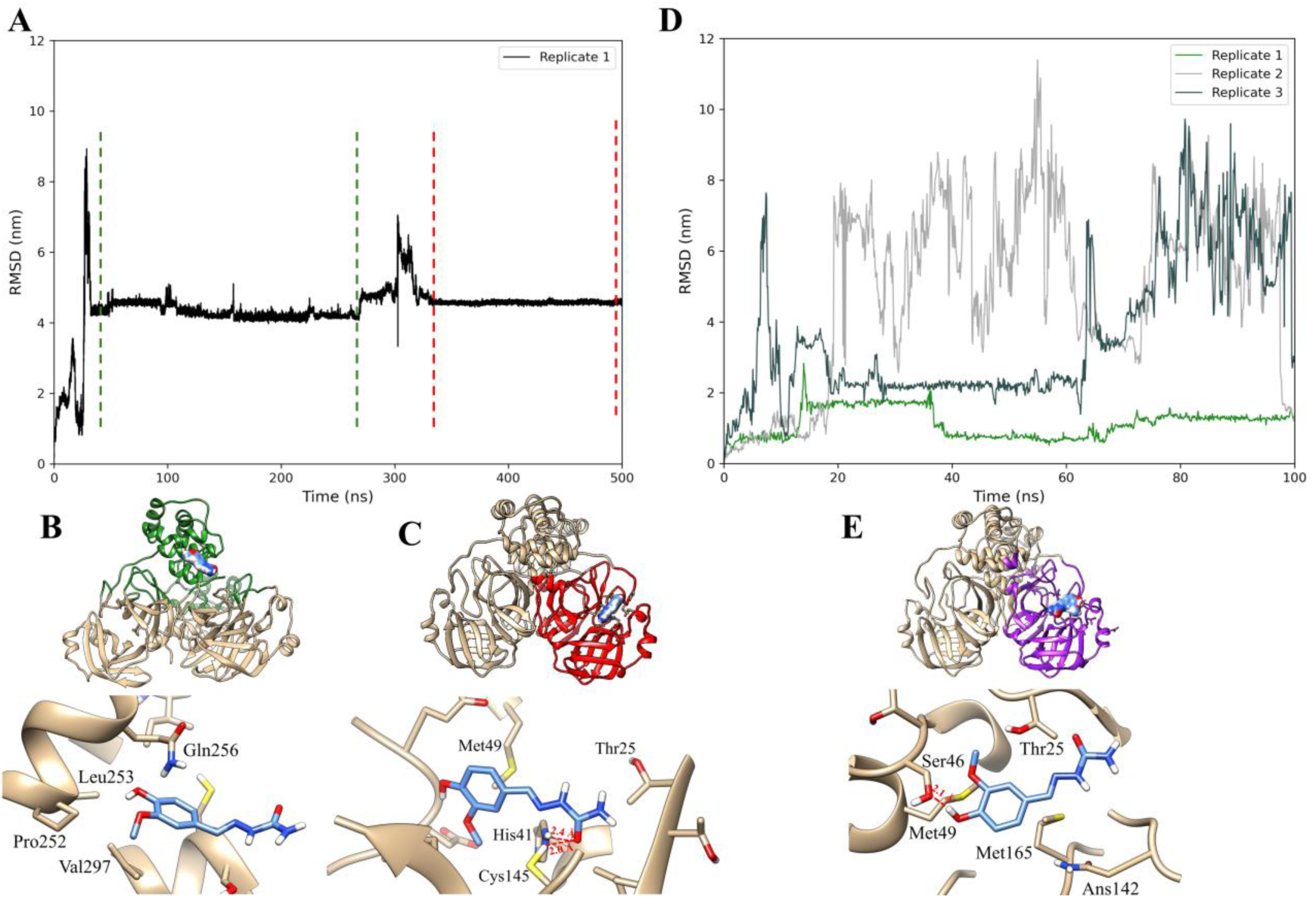
Molecular dynamics simulations suggest compound **25** as a potential competitive inhibitor of SARS-CoV-2 M^pro^. (**A**) RMSD (nm) of the top-ranked binding pose from the DockThor virtual screening mode, used as the initial pose over a 500 ns simulation. Representative structures of the most populated clusters of compound **25** observed during the simulation at the (B) allosteric site, shown as green ribbons, and (C) at the active site, shown as red ribbons. (**D**) RMSD profiles from three independent 100 ns simulations initiated from the predominant active site-bound conformation. **(E)** Representative binding pose of compound **25** from these replicates, highlighting key protein–ligand interactions. Dashed red lines represent potential hydrogen bonds. Figure generated using UCSF Chimera software.

Clustering analysis of the final 200 ns of the simulation indicated that the most representative cluster of compound **25** exhibited favorable interactions within the M^pro^ active site. The carbonyl oxygen of the semicarbazone moiety is oriented to potentially interact with His41 or Cys145 (Figure 3C), the residues comprising the catalytic dyad. Although the carbonyl group lies proximal to Cys145, it is unlikely to serve as an electrophilic center for nucleophilic attack due to resonance stabilization by the adjacent nitrogen atoms, which reduces the electrophilicity of the carbonyl carbon. Nevertheless, hydrogen bonding involving His41 and Cys145, combined with hydrophobic contacts with Met49, contributes to stabilizing compound **25** within the active site. On the other hand, the most populated cluster at the allosteric site is consistent with the low stability of compound **25** in this region, as observed between 50 and 260 ns of the simulation, with an average RMSD of 4.74 Å. Despite a potential hydrogen bond between its hydroxyl group and Gln256 (Figure 3B), the polar region of the ligand remains largely solvent-exposed, likely contributing to its diminished stability at this site. This is consistent with the predominantly hydrophobic nature of allosteric site 1, which preferentially accommodates less polar compounds.

Given the promising binding potential of compound **25** at the SARS-CoV-2 M^pro^ active site, the most populated cluster representing this interaction was selected as the starting conformation for additional MD simulations. RMSD analysis from three independent 100 ns replicates revealed that compound **25** does not maintain stable binding throughout all simulations (Figure 3D). However, in replicate 1, the ligand partially dissociated from the active site around 20 ns and subsequently reoriented within the binding pocket around 40 ns, suggesting dynamic interactions consistent with a reversible binding mode (Figure 3E). The most representative cluster from replicate 1 revealed that compound **25** adopted a shifted position within the M^pro^ active site relative to its initial position. Although it no longer directly engages the catalytic dyad (Cys145 and His41), the ligand remains within the substrate-binding pocket, which includes Ser46, Gln189, Thr190, Ala191, Pro168, Glu166, Leu141, and Asn142. This region encompasses the S1, S1′, S2, and S4 subsites, which are highly conserved among coronavirus main proteases. Specifically, the phenolic hydroxy group of compound **25** formed a potential hydrogen bond with Ser46 and engaged in hydrophobic interactions with Met49 and Met165 (Figure 3E). These residues have frequently been implicated in the binding of other known M^pro^ inhibitors.^64,65^ Therefore, although partial instability was observed in two of the simulation replicates, the reproducible interaction pattern of compound **25** within the active site supports its potential as a competitive inhibitor. To further characterize M^pro^ inhibition by compound **25**, we experimentally investigated the mechanism of inhibition. Kinetic analysis based on the Michaelis-Menten model (Figure 4A) yielded a K_i_ of 87.4 μM for compound **25**. Lineweaver-Burk plot analysis revealed a competitive inhibitor profile, as evidenced by an increase in the apparent Kₘ with increasing inhibitor concentration, whereas Vₘₐₓ remained unchanged (Figure 4B). These results are consistent with compound **25** acting as a reversible active-site-directed inhibitor.

**Figure 4.**
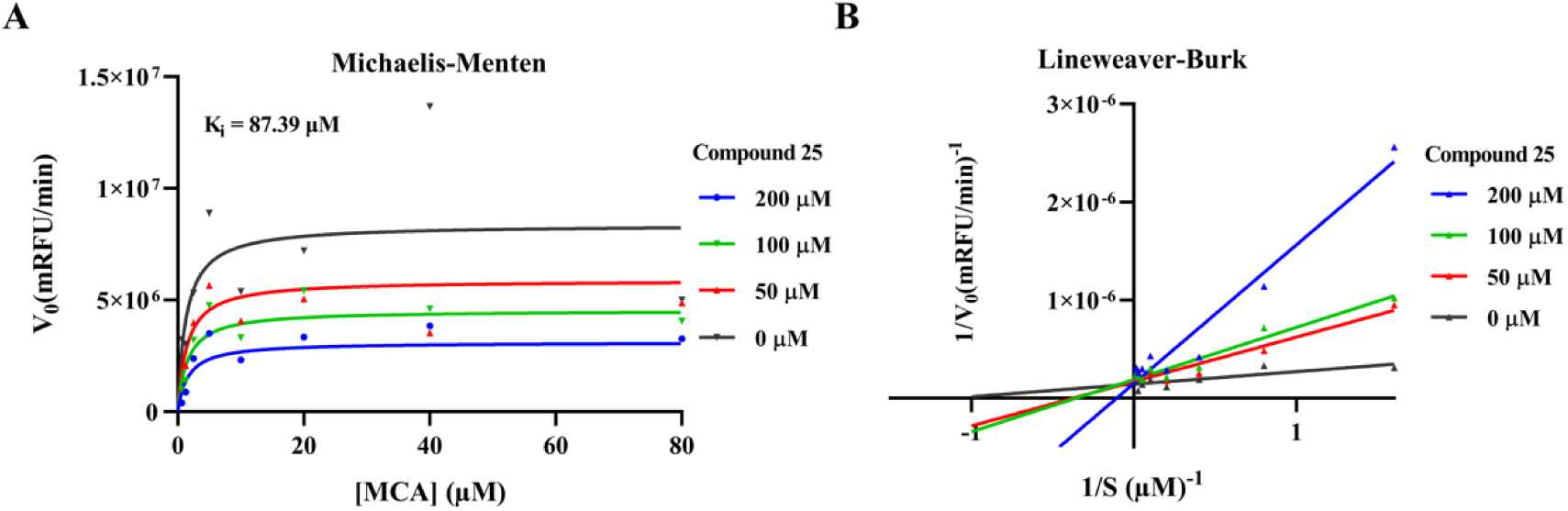
Determination of IC_50_ and inhibition mechanism of SARS-CoV-2 M^pro^ by compound **25**. (A) Kinetic curves for SARS-CoV-2 M^pro^ inhibition in the presence of three concentrations of compound **25** (50–200 µM) and increasing MCA-AVLQSGFR-Lys(Dnp)-Lys-NH2 substrate concentrations (ranging from 0.62 to 80 µM) fitted to the Michaelis-Menten model. (B) Lineweaver-Burk plot showing the effect of three concentrations of **25** on enzyme kinetics (SARS-CoV-2 M^pro^ at 40 nM).

### 2.4 SAR and discovery of additional SARS-CoV-2 M^pro^ inhibitors

To investigate the structure-activity relationship (SAR) within this chemical series, we selected 11 analogs of compound **25** (compounds **42-51**) from an *in-house* chemical library. Each compound was evaluated for its inhibitory activity against SARS-CoV-2 M^pro^ at a concentration of 100 μM, except for compound **46**, which was tested at 50 μM owing to limited solubility (Table 1). This set of analogs enabled the exploration of structural variations, including modifications to the phenyl ring and the substitution of semicarbazone moieties with thiosemicarbazone counterparts.

**Table 1.**
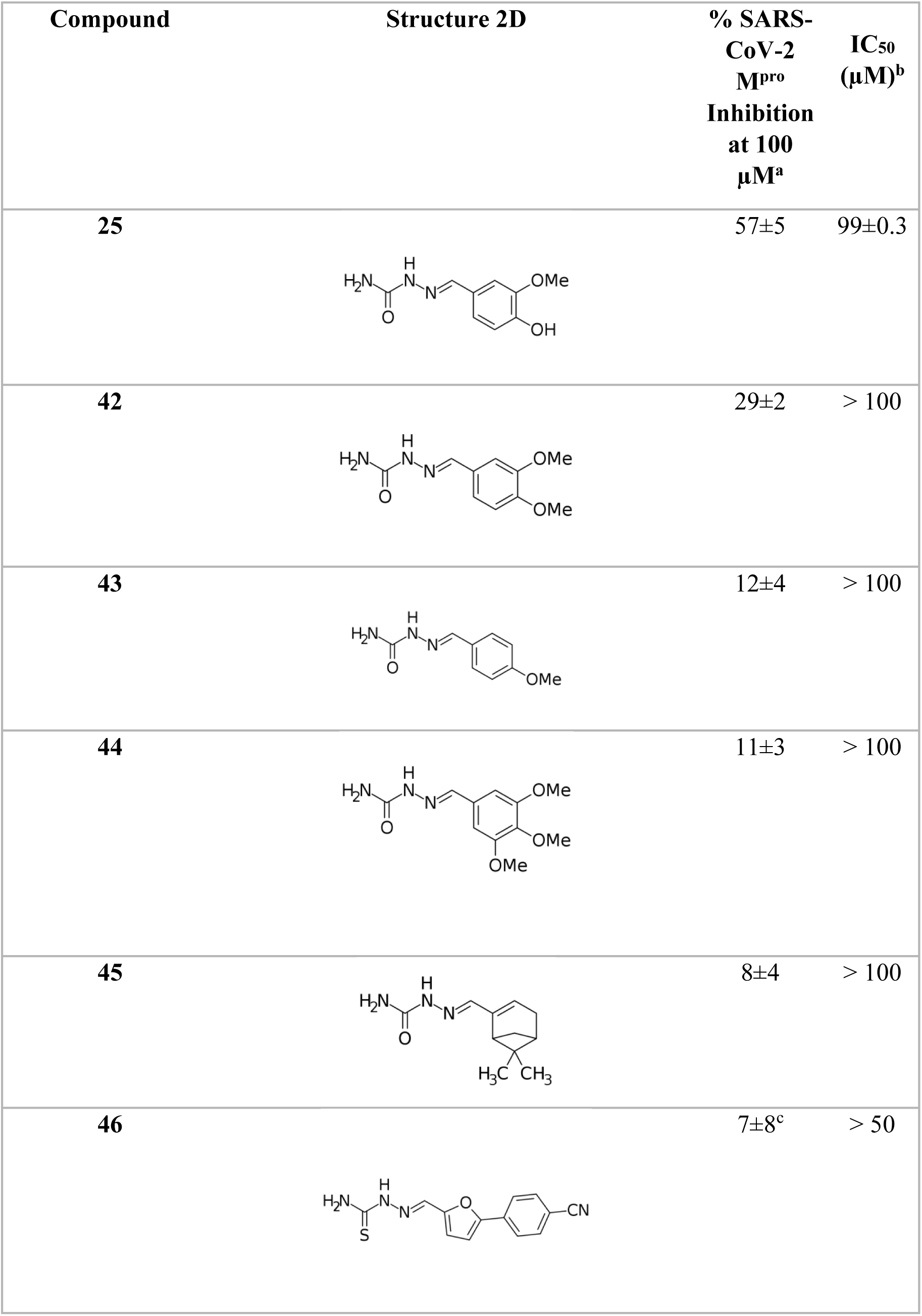
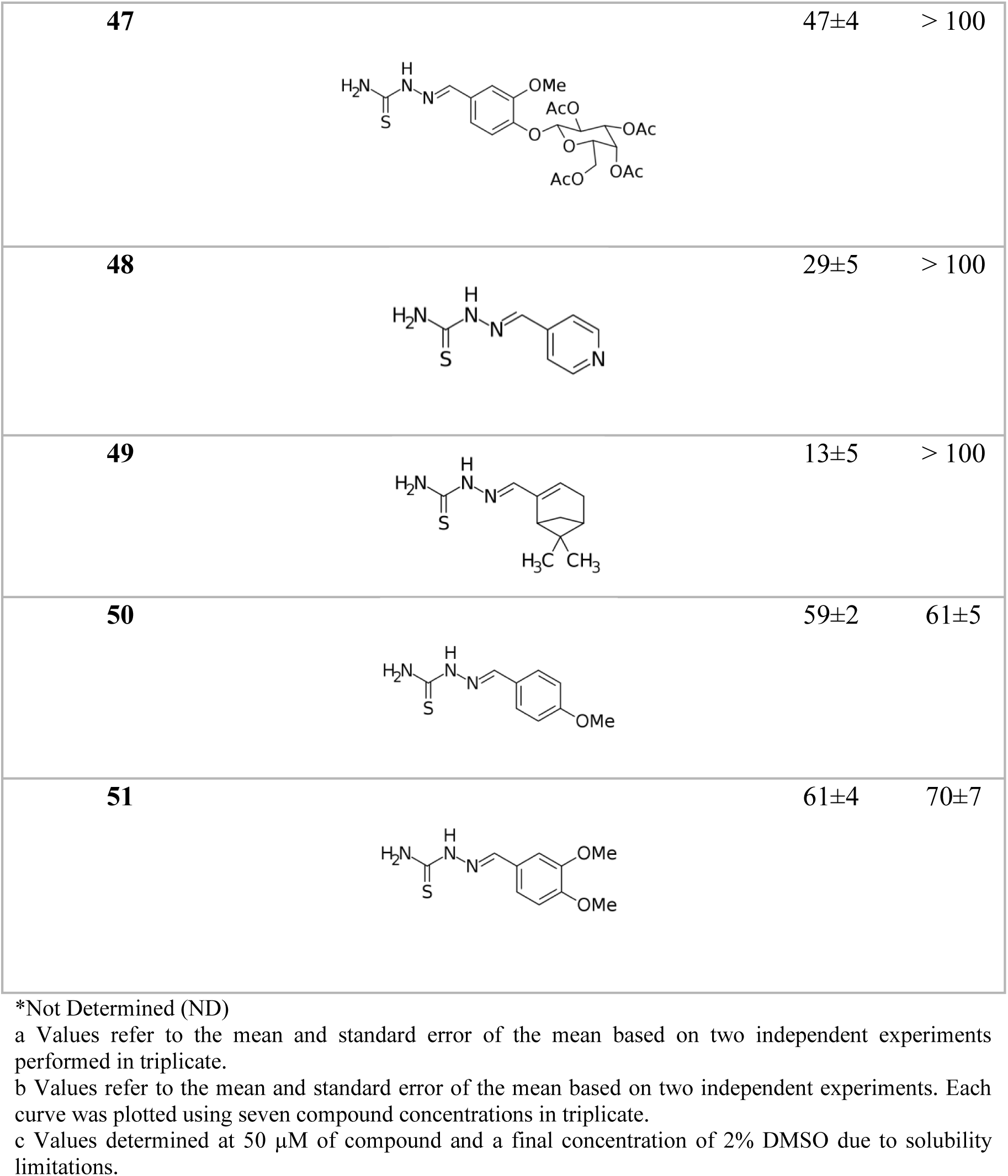
Inhibitory Effects and IC_50_ values for compound 25 analogs against SARS-CoV-2 M^pro^.

| Compound | Structure 2D | % SARS-CoV-2 M<sup>pro</sup> Inhibition at 100 μM<sup>a</sup> | IC<sub>50</sub> (μM)<sup>b</sup> |
| --- | --- | --- | --- |
| 25 | | 57±5 | 99±0.3 |
| 42 | | 29±2 | > 100 |
| 43 | | 12±4 | > 100 |
| 44 | | 11±3 | > 100 |
| 45 | | 8±4 | > 100 |
| 46 | | 7±8<sup>c</sup> | > 50 |
| **47** |  | 47±4 | > 100 |
| **48** |  | 29±5 | > 100 |
| **49** |  | 13±5 | > 100 |
| **50** |  | 59±2 | 61±5 |
| **51** |  | 61±4 | 70±7 |

\*Not Determined (ND)
a Values refer to the mean and standard error of the mean based on two independent experiments performed in triplicate.
b Values refer to the mean and standard error of the mean based on two independent experiments. Each curve was plotted using seven compound concentrations in triplicate.
c Values determined at 50 µM of compound and a final concentration of 2% DMSO due to solubility limitations.

Overall, modifications to the substitution pattern on the phenyl ring of compound **25** led to reduced SARS-CoV-2 M^pro^ inhibition. This trend was evident upon methylation or removal of the phenolic hydroxyl group (analog **42**, bearing a 4,5-dimethoxyphenyl: IC_50_ > 100 µM; analog **43**, bearing a 4-methoxyphenyl: IC_50_ > 100 µM). Similarly, compound **44**, bearing a 3,4,5-trimethoxyphenyl group, demonstrated 11 ± 3% inhibition at 100 µM. These findings highlight the importance of the hydroxyl substituent in **25,** consistent with our molecular dynamics simulations, which revealed a stable binding mode in which this moiety forms a hydrogen bond with the side chain of SARS-CoV-2 M^pro^ residue Ser46. In addition, replacing the phenyl with a bicyclo-3.1.1-heptane ring system resulted in loss of activity, as observed for analog **45** (IC_50_ > 100 µM).

We also investigated thiosemicarbazone analogs, which generally exhibited higher inhibition values than their semicarbazone counterparts, as commonly reported in the literature.^66–70^ Thiosemicarbazones **50** (bearing a 4-methoxyphenyl, 59±2% inhibition) and **51** (bearing a 3,4-dimethoxyphenyl, 61±4% inhibition) showed increased inhibitory activity compared to the semicarbazone analogs **43** (12 ± 4% inhibition) and **42** (29 ± 2% inhibition), respectively (Table 1). These compounds showed IC_50_ values of 61±5 for **50** and 70±7 μM for **51** (Figure 5A-B). Mechanism of inhibition analysis for compound **50** revealed a competitive inhibitor profile, comparable to that of compound **25**, with a *K*_i_ of 18.2 μM (Figure 5C-D).

**Figure 5.**
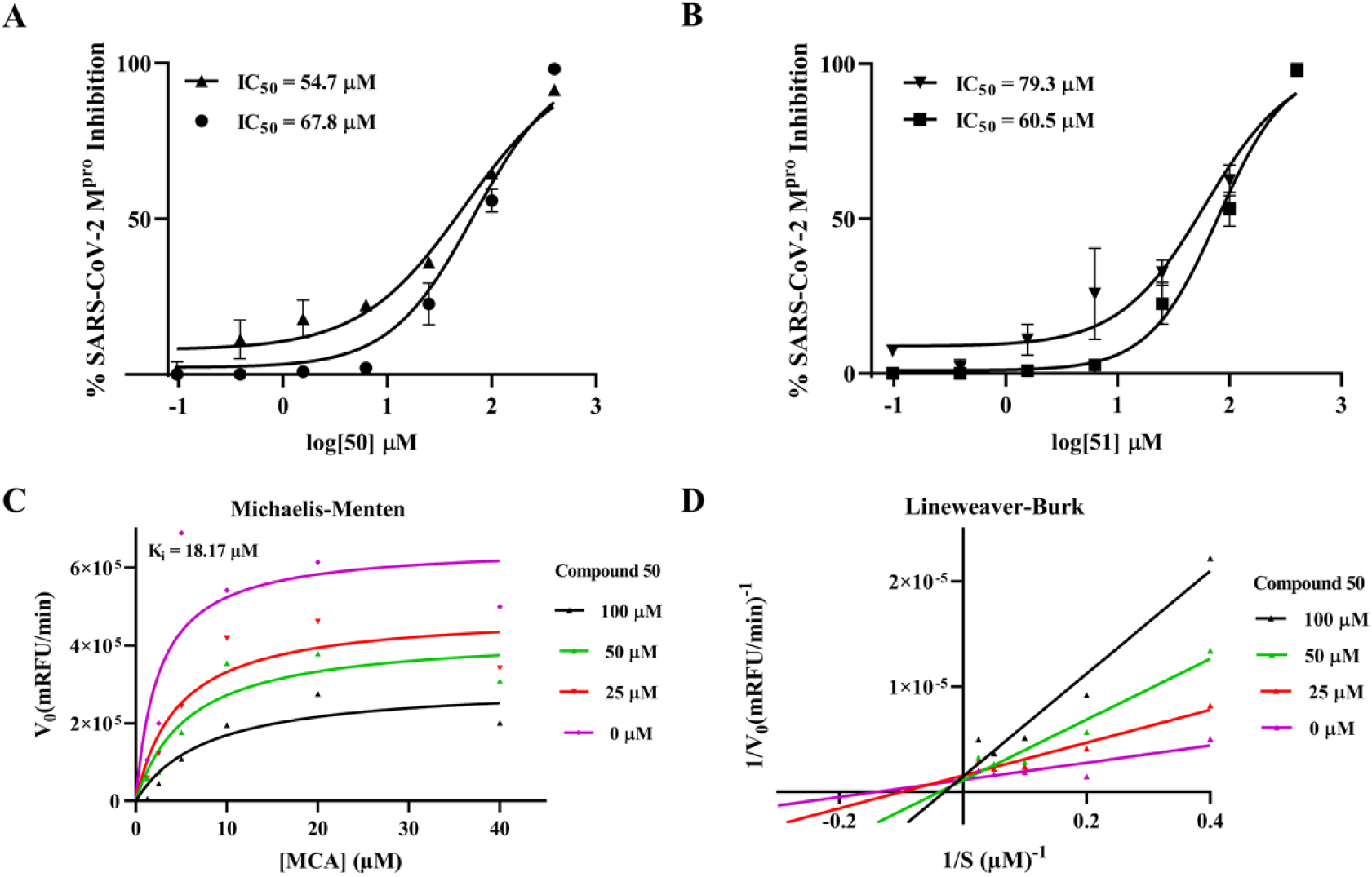
Determination of the IC_50_ of compounds **50** and **51** and the mechanism of SARS-CoV-2 M^pro^ inhibition by compound **50.** (A) Concentration-response curves for SARS-CoV-2 M^pro^ inhibition by **50** (concentration range: 0.097–400 µM). (B) Concentration-response curves for SARS-CoV-2 M^pro^ inhibition by **51** (concentration range: 0.097–400 µM). Each IC_50_ curve represents an independent experiment performed following preincubation with the enzyme, with triplicates measurements for each compound concentration. (C) Kinetic curves for SARS-CoV-2 M^pro^ inhibition in the presence of three concentrations of compound **50** (25–100 µM) and increasing MCA-AVLQSGFR-Lys(Dnp)-Lys-NH2 substrate concentrations (ranging from 1.25 to 40 µM) fitted to the Michaelis-Menten model. (D) Lineweaver-Burk plot showing the effect of three concentrations of **50** on enzyme kinetics (SARS-CoV-2 M^pro^ at 40 nM).

The enhanced inhibitory activity of thiosemicarbazones is frequently attributed to their increased electrophilicity and their ability to inhibit cysteine proteases by these compounds, through covalent binding.^67,69,71^ Thiosemicarbazones have been widely explored as inhibitors of cysteine proteases, including SARS-CoV-2 M^pro^.^69,71^ In a recent study by our group, we identified four thiosemicarbazones as covalent inhibitors of SARS-CoV-2 M^pro^, including one structural analog of the hits reported here.^69^ However, the absence of time-dependence observed for inhibition by compounds **50** and **51** (Table 2) suggests that these molecules interact with M^pro^ through noncovalent interactions, distinguishing them from previously reported inhibitors. Additionally, most previously described thiosemicarbazone inhibitors present a more complex scaffold than the one reported here.^69^

**Table 2.**
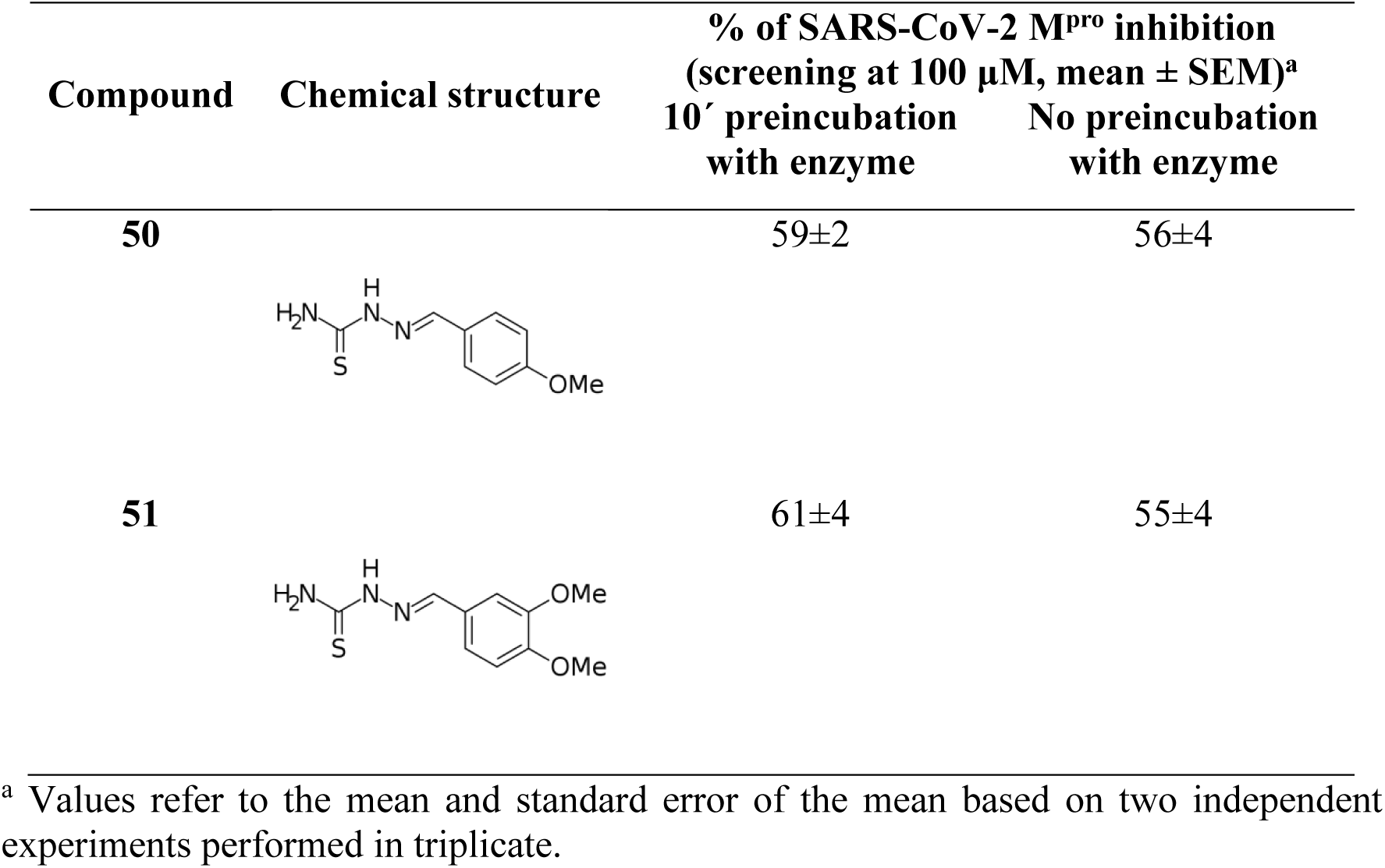
Influence of enzyme preincubation in SARS-CoV-2 M^pro^ inhibition by compounds 50 and 51.

| Compound | Chemical structure | % of SARS-CoV-2 M<sup>pro</sup> inhibition<br>(screening at 100 μM, mean ± SEM)<sup>a</sup> | |
| --- | --- | --- | --- |
|  |  | 10' preincubation<br>with enzyme | No preincubation<br>with enzyme |
| **50** | | 59±2 | 56±4 |
| **51** | | 61±4 | 55±4 |

<sup>a</sup> Values refer to the mean and standard error of the mean based on two independent experiments performed in triplicate.

Several compounds evaluated in the present study were also characterized in an earlier study targeting parasite cysteine proteases, including *Trypanosoma brucei* cathepsin L (*Tbr*CatL, also known as rhodesain), cruzain, and cathepsin B1 from *Schistosoma mansoni* (SmCB1).^66^ For instance, analog **47** (47 ± 4% inhibition against SARS-CoV-2 M^pro^), bearing an acetylated glycoside moiety, inhibited *Tbr*CatL (IC_50_ = 1.2 ± 1 μM) and cruzain (IC_50_ = 37.7 ± 10 μM). Analog **51** also exhibited low micromolar potency against *Tbr*CatL (IC_50_ = 6.2 μM), cruzain (IC_50_ = 9.0 μM), and SmCB1 (IC_50_ = 14.6 μM). In contrast, analog **50** was inactive against these three parasitic proteases, suggesting a degree of selectivity and supporting the specificity of its inhibitory activity against SARS-CoV-2 M^pro^.

## 3. Conclusion

In this study, we employed a structure-based virtual screening strategy targeting an allosteric site of SARS-CoV-2 M^pro^, leading to the identification of compound **25** as a novel inhibitor. Molecular dynamics simulations revealed an unexpected but stable relocation of compound **25** to the SARS-CoV-2 M^pro^ active site, where it exhibited key interactions consistent with competitive inhibition. Biochemical assays confirmed its inhibitory activity with a *K*_i_ value of 87.4 μM. Structure–activity relationship studies of analogs revealed the importance of the phenolic hydroxyl group and that conversion to thiosemicarbazones significantly enhanced SARS-CoV-2 M^pro^ inhibition, as exemplified by analog **50** (*K*_i_ = 18.2 μM). Despite prior evidence that thiosemicarbazones can act as covalent inhibitors of cysteine proteases, our data indicate that compounds **25**, **50**, and **51** act via a noncovalent, reversible mechanism. Importantly, the observed selectivity of analog **50** for SARS-CoV-2 M^pro^ over parasite cysteine proteases supports its specificity.

## 4. Methods

### 4.1 Redocking and Cross-docking Analysis

The DockThor server ^61^ (freely available at https://www.dockthor.lncc.br) and AutoDock Vina^60^ were used to perform redocking and cross-docking analyses. Redocking analyses were performed with the crystallographic structures of the complexes with the ligands ifenprodil from allosteric site 1 and AT-7519 from allosteric site 2 (PDB: 7AQI and 7AGA, respectively). The crystallographic ligands used for cross-docking analyses on allosteric site 1 were extracted from the PDB: 7AMJ (PD-168568), 7ABU (RS-102895), 7APH (tofogliflozin), and 7AXM (pelitinib). Protein and ligand structures were downloaded from the PDB^72^. We used an *in-house* script to calculate the RMSD between docking results and the experimentally determined binding mode based on equation (1)^73^:

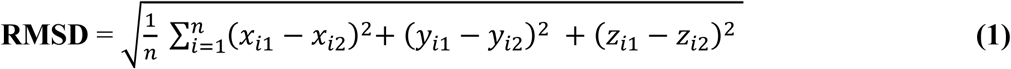

Where *x_i_*, *y_i_*, and *z_i_* are the coordinates of a pair of conformations 1 and 2 of a given molecule, and *n* is the number of atoms of the same molecule.

### 4.2 Ligand Preparation

The 2D structure of ligands for redocking and cross-docking was drawn in the Marvin Sketch 16.10.3 software (Chemaxon, 2015). The most abundant protonation state was predicted at a pH of 7.5. Next, using Avogadro software (Cheminformatics, 2012), we performed geometry and energy minimization employing the General AMBER Force Field (GAFF) with 1000 steps of the Steepest Descent algorithm, and convergence of 10^e-9^.

### 4.3 Protein Preparation

The structures 7AQI and 7AGA were downloaded from the PDB website. We used the H++ platform (http://newbiophysics.cs.vt.edu/H++/) to remove alternative conformations of residues with lower occupancy, water molecules, add missing hydrogen atoms, and define protonation states of amino acid residues at pH 7.5.

### 4.4 Grid Preparation

We used AutoDock Tools 1.5.6 (ADT) software to prepare the grid. For each allosteric site, docking grids were constructed based on the center of mass of the respective ligand’s coordinates (ifenprodil, from PDB: 7AQI, and AT-7519, from PDB: 7AGA), using the Discovery Studio Visualizer software (BIOVIA, 2020). They included the residues that interacted with their respective ligands. The spacing of the box was 0.375 Å. For the allosteric site 1, the grid size has the following coordinates: x = 16 Å, y = 16 Å, and z = 18 Å. The grid center coordinates were x = 4.910, y = 3.475, and z = -21.832. For allosteric site 2, the grid size was designed using the following coordinates: x = 20 Å, y = 20 Å, and z = 20 Å, with the grid center coordinates x = 17.088, y = 3.4133, z = -6.133.

### 4.5 Virtual screening libraries

Two libraries were used: one with 1,176 compounds (BraCoLi) and another with 884 compounds (ENSJ). BraCoLi is a library available openly at https://www.farmacia.ufmg.br/qf/downloads/^59^. BraCoLi ligands were downloaded in the .mol2 format. They had been previously prepared using the following protocol: 2D chemical structures were generated in Marvin Sketch 16.10.3 (Chemaxon, 2015), converted to 3D format, and their conformational energy was minimized using Discovery Studio Visualizer (BIOVIA, 2020). Additionally, any missing hydrogen atoms were added to the structures. The most stable conformers were generated by OMEGA 2.5.1.4. Ionization states at physiological pH (7.4) were corrected using the Fixpka 23 software implemented in QUACPAC 1.6.3.1 (OpenEye Scientific Software, 2016), in which the total energy was minimized using the MMFF94 force field. The 3D structure dataset was made available in SDF and Mol2 file formats. ENSJ is an *in-house* library, previously published and prepared with the following protocol: ligand 3D structures were generated from SMILES with LigPrep (version 46013), using Epik to predict their protonation at pH 7.0±2, and generating tautomers and diastereoisomers.^5^ The OPLS3e force field was employed for structure minimization.

### 4.6 Virtual Screening

The chemical libraries were docked against the allosteric site 1 of SARS-CoV-2 M^pro^ using the same grid box (center of coordinates and size) and protein file prepared for crossdocking experiments, using DockThor. For virtual screening, 12 docking runs were performed, with 500,000 evaluations by the genetic algorithm per docking run, a population of 750 individuals, and a maximum of three cluster leaders on each docking run. DockThor is a program that performs a non-covalent-docking simulation, utilizing the MMFF94S49 force field to define the atom types and partial charges of the protein and ligand files.^13^ The scoring function “Total Energy (Etotal)”, employed to select the binding mode for each ligand, is physics-based and composed of the terms: torsional ligand, van der Waals (constant buffer δ = 0.35), electrostatic potentials between the protein-ligand pairs of atoms, the intermolecular interaction energy (calculated by the sum of the electrostatic and van der Waals powers between 1-4 pairs of atoms). The scoring function DockTScore, employed for ranking compounds, is empirical and explicitly accounts for interaction physics-based terms that contribute to the binding free energy. DockTScore considers the following terms: intermolecular interaction terms, lipophilic protein-ligand interaction term, torsional entropy term, polar solvation term, and favorable nonpolar solvation term. DockThor-VS web server uses a multiple solution steady-state genetic algorithm based on phenotypic clustering as the search method. We analyzed the best-ranking pose generated for each compound for visual analysis and ligand selection using the ViewDock tool of the UCSF Chimera 1.16 software (UCSF Resource for Biocomputing, Visualization, and Informatics, 2004). The compounds were organized based on the number of hydrogen bond interactions in the protein-ligand complex. Each ligand was visually inspected, considering complementarity within the site, observing the number of protein-ligand interactions and the residues involved in these interactions, and the number of polar groups involved (and those not involved) in interactions with the protein.

### 4.7 Chemistry

Compound **26** was purchased from Sigma (USA) and tested without further purification. The remaining compounds were synthesized and characterized by our group. In most cases, the data has been previously published, as detailed below. In the case of compounds synthesized during this study, we describe their synthesis and characterization and provide the corresponding spectral data in the Supplementary Information 2 file.

The synthesis and characterization of the following compounds has been previously reported: compounds **2**, **7** and **29** ^74^; compound **6** ^69^; compounds **3**, **4**, **8**, **16**, **30** ^75^; compound **9** ^76^; compound **11** ^77^; compounds **12** and **46** ^78^; compound **13** ^79^; compounds **14**, **15**, **17** ^80^; compounds **10**, **18** and **19** ^81^; compound **20** ^82^; compound **22** ^5^; compounds **25**, **42**, **43**, **44**, **45**, **49**, **50** and **51** ^70^; compound **24** ^83^; compound **31** ^84^; compound **33** ^85^; compound **34** ^86^; compounds **35**, **37** and **40** ^87^; compounds **36** and **38** ^88^; compound **39** ^89^; compounds **47** and **48** ^66^.

#### Synthesis of compound **1**

The synthesis and characterization of compound **1** was carried out from: 2,5-anhydro-1,6-bis[(3,4-dichlorobenzoyl)amino]-1,6-dideoxy-D-mannitol^90^. In a 25 mL round-bottom flask 1,6-diamino-2,5-anhydro-1,6-didexoxy-D-mannitol^91^ (60 mg, 0.37 mmol) was dissolved in 2.0 mL of distilled water. Then, sodium bicarbonate (78 mg, 0.93 mmol) and 0.5 mL of THF were added. Separately, 3,4-dichlorobenzoyl chloride (171 mg, 0.81 mmol) was solubilized in 1.3 mL of THF, and this solution was added to the reaction mixture dropwise in an ice bath. The system was kept under magnetic stirring at room temperature for 20 hours, when the consumption of the starting material was verified by TLC (eluent: ethyl acetate, developer: UV light and ninhydrin solution). The THF was then removed on a rotary evaporator, and to the remaining liquid was added 15 mL of saturated sodium bicarbonate solution. Extraction was carried out with ethyl acetate (3 x 20 mL), and the organic phase was washed with distilled water (2 x 20 mL) and saturated sodium chloride solution (1 x 20 mL), dried with anhydrous sodium sulfate, filtered, and concentrated to dryness on a rotary evaporator. The crude material obtained was recrystallized from chloroform affording 100 mg of a white solid corresponding to the product (53% yield). Melting point: 175-177 °C; [α]D +40 (c 0,5, MeOH). 1H NMR (400 MHz, DMSO-d6), δ(ppm): 8.68 (t, J 5.5 Hz, 2H), 8.06 (d, J 1.8 Hz, 2H), 7.81 (dd, J 8.4/1.8 Hz, 2H), 7.72 (d, J 8.4 Hz, 2H), 5.30 (d, J 3.3 Hz, 2H), 3.99–3.91 (m, 2 H), 3.86 – 3.79 (m, 2), 3.50-3.37 (m, 4H). 13C NMR (100 MHz, DMSO-d6), δ(ppm): 164.1, 134.8, 133.9, 131.1,130.6, 129.2, 127.5, 82.5, 79.4, 41.2.

#### Synthesis of compound **5**

Prepared according to Lavorato et al.,^74^ except that p-toluenesulfonyl chloride was used in place of methanesulfonyl chloride. Yield: 95%. Melting point: 164-167 °C; 1H NMR (200 MHz, DMSO-d6), δ (ppm): 8.18 (d, J 9 Hz, 4H), 7.79 (d, J 8.2 Hz, 4H), 7.40 (d, J 8.2 Hz, 2H), 7.02 (d, J 9 Hz, 4H), 5.20 (m, 1H), 4.46 (d, J 4,2 Hz, 4H), 2.38 (s, 3H). 13C NMR (50 MHz, DMSO-d6), δ (ppm): 162.8, 145.2, 141.2 132.9, 130.0, 127.8, 125.8, 115.1, 78.1, 67.2, 21.1.

#### Synthesis of compounds **21**, **23**, **27**, and **28**

To a cooled to 0 °C solution of Nα-benzyloxycarbonyl-β-amino-L-alanine (0.84 mmol) in NaOH 1 mol/L solution (2 mL) and THF (5 mL) was added dropwise the appropriate cinnamic acid chloride (0.68 mmol) previously dissolved in THF (5 mL). The reaction mixture was stirred for 15h, maintaining pH at 10–12 with NaOH 1 mol/L water solution. The course of the reaction was followed by TLC. After completion, THF was evaporated and water and HCl 1 mol/L solution were added to pH 1. EtOAc was added and the organic phase was washed with H2O. The organic phase, dried with MgSO4, was evaporated and the crude purified by flash chromatography on silica gel (cyclohexane/ EtOAc 1:1; cyclohexane/ EtOAc 3:7) to afford the amides which were considered pure enough by TLC analysis to be used in the next step. To a solution of corresponding amide (0.41-0.57 mmol) in acetic acid (5 mL) cooled to 0 °C, 1 mL of 33% HBr in acetic acid (0.57 mmol) was added and then the reaction mixture was stirred for 2-4 hours. The mixture was concentrated to a small volume diluted with water (15 mL) and extracted with ether (3x15 mL). The water phase was evaporated. The solid obtained was washed with acetone to give compounds 21, 23, 27 and 28.

#### Compound **21**

0.108 g (55% yield). Melting point: 246.4-248.5 °C. [α]_D_ ^23^ -4.9 (*c* 0.2, TFA/H_2_O 5%). HRMS (ESI^+^) *m/z*: [M+H]^+^ calculated 313.0188, found 313.0186. ^1^H NMR (400 MHz, D_2_O/TFA) *δ* (ppm): 6.23 (d, *J* 16 Hz, 1H), 6.21 (d, *J* 8.4 Hz, 2H), 6.12 (d, *J* 8.4 Hz, 2H), 5.33 (d, *J* 16 Hz, 1H), 3.20 (dd, *J* 6.4/4.0 Hz 1H), 2.84 (dd, *J* 16/4 Hz, 1H), 2.72 (dd, *J* 16/6.4 Hz, 1H). ^13^C NMR (100 MHz, D_2_O/TFA) *δ* (ppm): 173.4, 172.8, 145.0, 136.4, 135.1, 132.7, 127.2, 122.5, 55.8, 42.7.

#### Compound **23**

0.180 g (78% yield). Melting point: 220.8-223.9 °C. [α]_D_ ^24^ -1.67 (*c* 0.3, TFA/H_2_O 5%). HRMS (ESI^+^) *m/z*: [M+H]^+^ calculated 293.1137, found 293.1137. ^1^H NMR (400 MHz, D_2_O/TFA) *δ* (ppm): 6.32 (d, *J* 8.4 Hz, 2H), 5.94 (d, *J* 16 Hz, 1H), 5.92 (d, *J* 8.4 Hz, 2H), 5.03 (d, *J* 16 Hz, 1H), 2.76 (dd, *J* 6/4 Hz 1H), 2.43 (dd, *J* 16/4 Hz, 1H), 2.30 (dd, *J* 16/6 Hz, 1H), 2.26 (s, 3H). ^13^C NMR (100 MHz, D_2_O/TFA) *δ* (ppm): 173.8, 173.0, 172.5, 145.5, 142.4, 134.3, 133.4, 131.5, 124.5, 57.4, 55.8, 43.2.

#### Compound **27**

0.110 g (56% yield). Melting point: 230.3-233.8 °C. [α]_D_ ^23^ -7.0 (*c* 0.21, TFA/H_2_O 5%). HRMS (ESI^+^) *m/z*: [M+H]^+^ calculated 269.0692, found 269.0692. ^1^H NMR (400 MHz, D_2_O/TFA) *δ* (ppm): 6.69 (d, *J* 15.6 Hz, 1H), 6.67 (d, *J* 8.4 Hz, 2H), 6.55 (d, *J* 8.4 Hz, 2H), 5.77 (d, *J* 15.6 Hz, 1H), 3.68 (dd, *J* 6/4 Hz, 1H), 3.29 (dd, *J* 16/4 Hz, 1H), 3.18 (dd, *J* 16/6 Hz, 1H). ^13^C NMR (100 MHz, D_2_O/TFA) *δ* (ppm): 172.6, 172.4, 144.0, 138.3, 135.6, 132.1, 131.7, 122.3, 56.1, 42.1.

#### Compound **28**

0.115 g (61% yield). Melting point: 184.6-189.5 °C. [α]_D_ ^23^ -4.2 (*c* 0.12, TFA/H_2_O 5%). HRMS (ESI^+^) *m/z*: [M+H]^+^ calculated 280.0933, found 280.0932. ^1^H NMR (400 MHz, D_2_O/TFA) *δ* (ppm): 7.14 (d, *J* 8 Hz, 1H), 7.04 (d, *J* 16 Hz, 1H), 6.81–6.76 (m, 2H), 6.68–6.66 (m, 1H), 5.67 (d, *J* 16 Hz, 1H), 3.54 (dd, *J* 6/4 Hz, 1H), 3.18 (dd, *J* 16/4 Hz, 1H), 3.06 (dd, *J* 16/6 Hz, 1H). ^13^C NMR (100 MHz, D_2_O/TFA) *δ* (ppm): 172.6, 172.1, 150.7, 141.0, 137.3, 133.5, 133.2, 132.2, 127.8, 126.8, 56.4, 42.3.

#### Synthesis of compound **32:**

To a 50 mL round-bottom flask was added O-propargyllapachol^76^ (0.30 mmol) dissolved in 1 mL of tetrahydrofuran, followed by β-D-glucoyranosyl azide^92^ (0.27 mmol), dissolved in 0.5 mL of tetrahydrofuran. Then, Cu(OAc)2.H2O 50% mol, dissolved in 0.5 mL of water and sodium ascorbate 60% mol, dissolved in 1 mL of water were added in a stepwise manner. The reaction mixture was stirred at room temperature for 4 h and monitored by TLC analysis. The tetrahydrofuran was removed by distillation at reduced pressure. The deacetylated glycosyl triazole 32 was added to Florisil and purified with silica column using ethyl acetate: MeOH/9:1 as mobile phase. 2-[1-(β-D-glucopyranosyl)-1,2,3-triazol-4-(methyl)oxy]-3-(3-methyl-2-butenyl)-1,4-naphthoquinone (32)

Yellow amorphous powder. Yield: 59%; Melting point: 118.5 – 120.7 °C. [α]D20 -15.8 (c 0.38; MeOH); IR ν_max_ 3282, 2912, 1652, 1593, 1092, 1047 cm-1; ^1^H NMR (400 MHz, DMSO-d6): δ 8.42 (s, 1H), 8.04-7.96 (m, 2H), 7.87-7.83 (m, 2H), 5.55 (d, 1H, J 9.2 Hz), 5.46 (s, 2H), 5.34 (d, 1H, J 6.4 Hz), 5.25 (d, 1H, J 4.6 Hz), 5.13 (d, 1H, J 5.6 Hz), 4.99 (t, 1H, J 7.2 Hz), 4.58 (t, 1H, J 5.2 Hz), 3.78-3.68 (m, 2H), 3.47-3.21 (m, 4H), 3.11 (d, 2H, J 7.2 Hz), 1.64 (s, 3H), 1.60 (s, 3H); 13C NMR (100 MHz, DMSO-d6) δ 184.6, 181.0, 156.2, 142.5, 134.1, 134.1, 133.7, 132.6, 131.3, 131.1, 125.9, 125.6, 123.9, 120.0, 87.4, 79.9, 76.9, 72.0, 69.5, 65.5, 60.6, 25.4, 22.6, 17.7; UPLC purity>99%, tR = 4.53 min; MS (ESI+) m/z calcd for C24H28N3O8 486.19, found 485.84 (M+H+).

#### Synthesis of compound **41**

In a 50 mL flask 0.44 g (1.50 mmol) of methyl 3-(*N*-cinnamoyl)amino-4-hydroxybenzoate was dissolved in a freshly prepared lithium hydroxide solution (0.06 g, 1.50 mmol) in 15 mL of distilled water under magnetic stirring for 10 minutes. To this solution was added 0.28 g (0.5 mmol) of 3,4,6-tri-*O*-acetyl-2-deoxy-2-(2-iodobenzoyl)amino-α-_D_-glucopyranosyl chloride dissolved in 10 mL of acetone. The reaction mixture was kept under magnetic stirring at room temperature. The reaction progress was monitored by TLC (eluent: hexane/ethyl acetate 1:1; developing agents: iodine vapor and 15% ethanolic sulfuric acid solution with heating). After the reaction was complete, the reaction mixture was concentrated under ventilation, the residue was dissolved in dichloromethane and a 6 mol/L hydrochloric acid solution was added. The mixture was filtered and the filtrate was extracted with 1 mol/L aqueous sodium hydroxide solution (4x 30 mL). The organic phase was washed with distilled water until pH 7, dried over anhydrous sodium sulfate and concentrated on a rotary evaporator under reduced pressure. The residue was purified by column chromatography (eluent: hexane/ethyl acetate 5.5:4.5) to yield 0.18 g (0.23 mmol) of the glycoside as a white solid (45.5% yield).

In a 250 mL three-tube round-bottom flask fitted with a reflux condenser, 0.25 g (0.31 mmol) of the glycoside and 30.0 mL of anhydrous benzene were added. The system was placed under reflux, nitrogen atmosphere, and magnetic stirring. A solution of 0.13 mL (0.15 g, 0.50 mmol) of Bu_3_SnH and AIBN (catalytic amount) dissolved in 10.0 mL of anhydrous benzene was added dropwise over a period of approximately 1 h. The system was maintained under reflux and magnetic stirring for 1 h after the addition was complete. The benzene was distilled under reduced pressure, and the residue obtained was pre-purified by column chromatography containing 10% KF (eluent:hexane/ethyl acetate 1:1). The fractions of interest were pooled, and further purification was carried out by flash column chromatography (eluent: dichloromethane/methanol 99.2:0.8). Compound **41** was isolated as white solid in 59.7% yield. Melting point: 286-288 °C; [α]_D_ 30.0 (*c* 0.2, CH_2_Cl_2_)

HRMS (ESI^+^) *m/z*: [M+H]^+^ calculated 689.2341, found 689.2103 ^1^H NMR (400 MHz, DMSO-d_6_) *δ* (ppm): 8.77 (d, *J* 9.6 Hz, 1H), 8.61 (s, 1), 8.46 (d, *J* 2.0 Hz, 1H), 7.73 (dd, *J* 8.8/2.4 Hz, 1H), 7.63 (d, *J* 7.6 Hz, 1H), 7.49 (td, 7.7/0.9 Hz, 1H), 7.38–7.26 (m, 6H), 7.20–7.16 (m, 2H), 5.35 (t, *J* 8.0 Hz, 1H), 5.24 (t, *J* 10.0 Hz, 1H), 5.08 (t, *J* 9.6 Hz, 1H), 4.93 (d, *J* 8.4 Hz, 1H), 4.54–4.47 (m, 1H), 4.31 (dd, *J* 12.4/4.8 Hz, 1H), 4.22 (dd, *J* 12.0/2.2 Hz, 1H), 4.14–4.10 (m, 1H), 3.82 (s, 3H), 3.58 (dd, *J* 14.0/8.8 Hz, 1H), 3.24 (dd, *J* 14.0/6.8, 1H), 2.09 (s, 3H), 2.05 (s, 3H), 2.04 (s, 3H) ^13C^ NMR (100 MHz, DMSO-d_6_) *δ* (ppm): 170.9, 170.6, 170.3, 170.1, 169.5, 165.6, 151.9, 139.3, 137.0, 134.8, 131.0, 129.0, 128.5, 126.3, 128.5, 128.0, 127.7, 127.3, 126.5, 125.2, 121.9, 117.1, 104.3, 71.8, 71.6, 68.1, 61.7, 53.5, 52.4, 47.2, 34.9, 20.7, 20.6, 20.5

### 4.8 Enzymatic assays with SARS-CoV-2 M^pro^

The expression and purification of recombinant SARS-CoV-2 M^pro^ were performed according to a previously established protocol.^23^ Enzymatic assays were performed in black flat-bottom 96-well plates using a Cytation 5 Cell Imaging fluorimeter (Biotek, Vermont, USA) with an excitation/emission value of 320/405 nm. SARS-CoV-2 M^pro^ activity was measured by cleavage of the fluorogenic substrate MCA-AVLQSGFR-Lys(Dnp)-Lys-NH2 trifluoroacetate (Sigma, US and Canada). The proteolysis was measured at 37 °C in 20 mM Tris-HCl buffer solution, pH 7.5, 0.5% DMSO, 1 mM DTT, 1 mM EDTA, 0.01% Tween-20 (all purchased from Sigma-Aldrich, San Louis, USA), 40 nM enzyme (SARS-CoV-2 M^pro^) and 10 µM substrate (K_m_ = 1.7 ± 0.8 µM), final plate volume of 100 µL. Initial screening was performed at concentrations of 25 μM to 100 µM of compounds after a 10-minute incubation with the enzyme, followed by the addition of a solution containing the substrate. The initial reaction velocities were calculated based on the first 5 min of reaction. Percentages of inhibition were calculated in comparison to the DMSO control. To determine the half-maximal inhibitory concentration (IC_50_) values, each curve was constructed using seven different concentrations of the compounds, and the IC_50_ values were determined by nonlinear regression analysis of the velocity vs. inhibitor. IC_50_ curves using GraphPad Prism 6 (GraphPad Prism, version 6, La Jolla, California, USA). The reported values are the average and standard deviation for at least two independent experiments in triplicate.

To determine the mechanism of enzyme inhibition by compound **25**, we evaluated inhibition at eight substrate concentrations, ranging from 80 to 0.62 μM. For compound **50**, we evaluated six concentrations ranging from 40 to 1.25 μM (2-fold serial dilutions), and 3 compound concentrations (one above the IC_50_, one similar to the IC_50_, and one below the IC_50_). The *K*_m_ and *V*_max_ values were determined by the nonlinear fit of the initial velocity results for each substrate concentration to the Michaelis−Menten or Mix-model equation^93^ using GraphPad Prism 6, using equation 2.

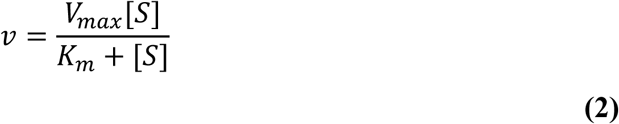

Where 𝑣 is the reaction rate, *V*_max_ is the maximum rate multiplied by the substrate concentration and divided by the sum of *K*_m_ plus [S].

*K*_i_ values were estimated based on equation 3:

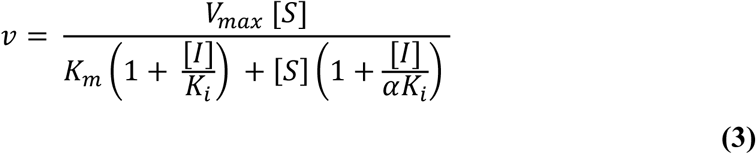

Where *K_i_* is the dissociation constant, one for the binary enzyme-inhibitor complex, and α*K*_i_ is the dissociation constant, another for the ternary complex: enzyme, substrate, and inhibitor (ESI).

### 4.9 Molecular Dynamics Simulations

All molecular dynamics (MD) simulations were performed using GROMACS version 2024.2.^94^ Ligand topologies were generated using the SwissParam server, which employs the Merck Molecular Force Field (MMFF) and provides parameters compatible with the CHARMM force field.^95^ Ligand coordinates were obtained from the top-ranked poses generated by molecular docking simulations conducted with the DockThor platform^64^, in both standard and virtual screening modes. The topology of the SARS-CoV-2 M^pro^ (PDB: 7AQI) was generated after preparation using the H++ platform (http://newbiophysics.cs.vt.edu/H++/), employing the CHARMM22 force field implemented in GROMACS. Each protein–ligand complex was solvated in a cubic box filled with TIP3P water molecules, ensuring a minimum distance of 10 Å between the protein and the box boundaries. System neutrality was achieved by the addition of counterions, and physiological ionic strength was maintained by supplementing with 0.15 M NaCl. Energy minimization was initially conducted using 5000 steps of the steepest descent algorithm, followed by 1000 steps of the conjugate gradient method. Minimization was carried out under constant volume (NVE) conditions, with electrostatic interactions computed using the Particle Mesh Ewald (PME) method. Following minimization, the systems underwent equilibration in two phases: NVT and NPT ensembles. Temperature was maintained at 300 °K using the V-rescale thermostat with a coupling constant of 0.1 ps. Pressure was maintained at 1 bar using the Parrinello–Rahman barostat. For the top-ranked docking pose from standard mode, three independent production and unrestrained simulations of 100 ns were carried out. The best-ranked pose pointed by virtual screening mode was subjected to a single replicate of 500 ns unrestrained MD analysis. GROMACS version 2024.2 was also employed to perform clustering analysis of simulation frames and to calculate RMSD values of the ligand, after fitting to the protein backbone.

## Supporting information

Spectral data

## 5. Acknowledgments

We would like to acknowledge OpenEye Scientific Software, Cadence Molecular Sciences for a free academic license, and the DockThor platform, AutoDock Vina, GROMACS. Avogadro, Marvin Sketch 16.10.3, Discovery Studio Visualizer, and UCSF Chimera for freely providing their software.

## 6. Declaration of generative AI and AI-assisted technologies in the writing process

During the preparation of this work the authors used ChatGPT and Grammarly in order to obtain suggestions to improve the clarity and grammar of text previously written by the authors. After using this tool/service, the authors reviewed and edited the content as needed and take full responsibility for the content of the published article. The text generated by the generative tool was not directly incorporated into the manuscript. It was critically analyzed by manual selection of expressions or suggestions for reorganization of specific sentences, which were then incorporated into the text written by the authors.

