## Supplementary Tables and Figures for "Discovery of Semicarbazone and Thiosemicarbazone Analogs as Competitive SARS-CoV-2 Virus Main Protease (M^pro^) Inhibitors"

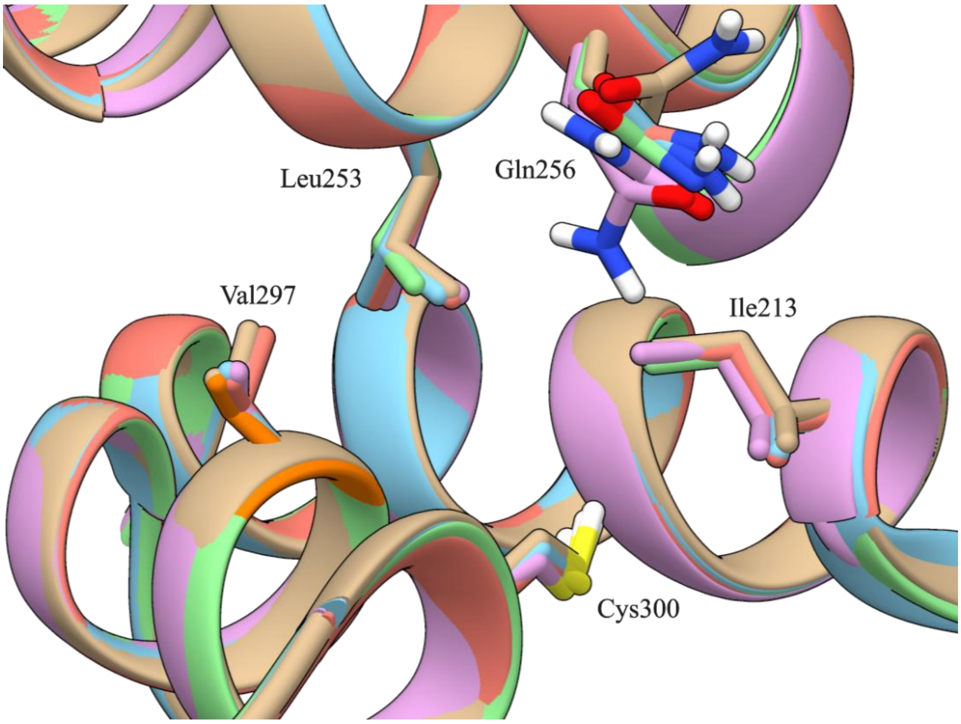


**Figure S1**. Conserved residues conformations in SARS-CoV-2 $M^{pro}$ allosteric site 1. Ile213, Leu253, Val297, Cys300, and Gln256 residues adopt similar conformations in the five crystallographic complexes superposed (PDBs: 7ABU, beige; 7AMJ, orange 7APH, pink; 7AQI, blue; 7AXM, green). SARS-CoV-2 $M^{pro}$ dimer is shown as cartoon and highlighted residues are depicted as sticks. Figure generated using UCSF Chimera software.

**Table S1 - Root mean square deviation (Å) for docking poses obtained in redocking and cross-docking performed by DockThor and AutoDock Vina.**

| Pose[c] | Ifenprodil[a] (site 1) | | Tofogliflozin (site 1) | | PD-168568[b] (site 1) | | Pelitinib (site 1) | | RS-102895[b] (site 1) | | AT-7519[a] (site 2) | |
|---|---|---|---|---|---|---|---|---|---|---|---|---|
| | Dock Thor | Auto Dock Vina | Dock Thor | Auto Dock Vina | Dock Thor | Auto Dock Vina | Dock Thor | Auto Dock Vina | Dock Thor | Auto Dock Vina | Dock Thor | Auto Dock Vina |
| **1** | 7.8 | 9.3 | 8.7 | 10.5 | 2.6 | 11.1 | 7.5 | 10.5 | 2.9 | 13.7 | 9.8 | 8.3 |
| **2** | 2.8 | 9.7 | 4.0 | 10.7 | 7.4 | 10.2 | 7.5 | 10.7 | 1.9 | 4.4 | 0.6 | 6.9 |
| **3** | 7.9 | 12.8 | 8.2 | 8.6 | 3.8 | 13.5 | 6.8 | 11.2 | 2.6 | 4.6 | 10.1 | 7.0 |
| **4** | 9.4 | 9.3 | 7.6 | 9.0 | 1.8 | 5.3 | 6.9 | 10.5 | 6.1 | 10.3 | 0.6 | 9.7 |
| **5** | 9.5 | 12.4 | 7.7 | 9.2 | 8.2 | 10.3 | 7.9 | 7.6 | 3.3 | 14.3 | 9.3 | 8.0 |
| **6** | 8.5 | 10.6 | 6.3 | 10.0 | 7.4 | 8.4 | 8.3 | 7.1 | 4.9 | 6.9 | 9.5 | 7.4 |
| **7** | 9.5 | 11.9 | 8.7 | 8.8 | 4.2 | 13.3 | 9.1 | 3.3 | 9.2 | 11.2 | 9.9 | 9.1 |
| **8** | 8.6 | 10.6 | 7.5 | 9.5 | 3.3 | 11.5 | 6.9 | 10.3 | 9.6 | 7.9 | 2.8 | 7.4 |
| **9** | 8.9 | 9.0 | | 8.9 | 3.9 | 5.8 | 7.1 | 7.4 | 9.7 | 6.9 | 5.3 | 7.8 |
| **10** | 8.4 | 8.9 | | 9.6 | 8.2 | 11.8 | 10.7 | 9.0 | 4.9 | 8.0 | 3.6 | 8.6 |

a Values refer to redocking experiments.

b Values refer to cross-docking experiments.

**Table S2. Docking scores (kcal/mol) for crystallographic ligand in redocking and cross-docking experiments performed with DockThor and AutoDock Vina.**

| | Ifenprodil[a] (site 1) | | Tofogliflozin (site 1) | | PD-168568[b] (site 1) | | Pelitinib (site 1) | | RS-102895[b] (site 1) | | AT-7519[a] (site 2) | |
|---|---|---|---|---|---|---|---|---|---|---|---|---|
| **Pose** | Dock Thor | Auto Dock Vina | Dock Thor | Auto Dock Vina | Dock Thor | Auto Dock Vina | Dock Thor | Auto Dock Vina | Dock Thor | Auto Dock Vina | Dock Thor | Auto Dock Vina |
| **1** | -7.6 | - 7.2 | -8.1 | -6.2 | -7.8 | -6.9 | -7.6 | -5.7 | -8.0 | -7.3 | -7.4 | -5.6 |
| **2** | -7.9 | -6.9 | -7.9 | -5.5 | -7.6 | -6.8 | -7.4 | -5.6 | -7.7 | -6.2 | -8.4 | -5.6 |
| **3** | -7.5 | -6.8 | -8.0 | -5.5 | -7.6 | -6.7 | -7.9 | -5.6 | -7.0 | -6.1 | -7.9 | -5.5 |
| **4** | -7.3 | -6.8 | -7.9 | -5.4 | -6.9 | -6.7 | -7.2 | -5.6 | -7.4 | -6.1 | -8.4 | -5.3 |
| **5** | -6.7 | -6.8 | -7.7 | -5.3 | -7.2 | -6.5 | -7.2 | -5.4 | -7.6 | -6.0 | -8.0 | -5.2 |
| **6** | -6.8 | -6.5 | -7.7 | -5.3 | -6.9 | -6.5 | -7.8 | -5.2 | -6.9 | -6.0 | -7.9 | -5.2 |
| **7** | -6.8 | -6.3 | -7.4 | -5.2 | -7.4 | -6.4 | -7.1 | -5.2 | -7.7 | -5.9 | -7.9 | -5.1 |
| **8** | -7.0 | -6.2 | -7.6 | -5.2 | -6.6 | -6.4 | -7.4 | -5.1 | -7.7 | -5.9 | -8.4 | -5.0 |
| **9** | -6.5 | -6.1 | | -5.1 | -7.7 | -6.4 | -7.2 | -4.9 | -7.6 | -5.9 | -7.7 | -5.0 |
| **10** | -7.6 | -6.1 | | -5.1 | -7.8 | -6.1 | -6.8 | -4.9 | -6.9 | -5.9 | -8.3 | -5.0 |

a Values refer to redocking experiments.

b Values refer to cross-docking experiments.

**Table S3.** Inhibitory effects of compounds against SARS-CoV-2 M$^{pro}$

| Compound ranking | Compound code | chemical structure | % of SARS-CoV-2 M$^{pro}$ inhibition (screening at 100 μM) |
|---|---|---|---|
| **11** | **1** | 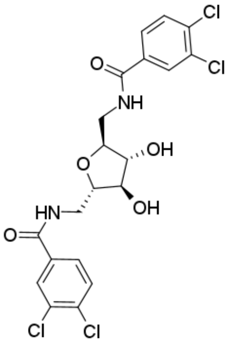 | **4±2[b]** |
| **27** | **2** | 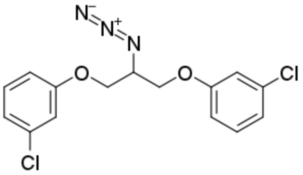 | **23±6[b]** |
| **29** | **3** | 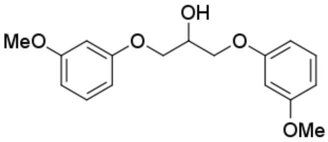 | **14±9** |
| **43** | **4** | 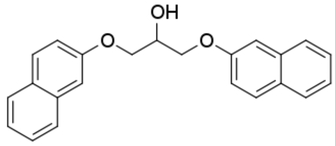 | **fluorescent compound[c]** |
| **45** | **5** | 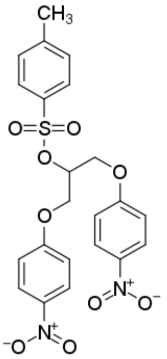 | **15±4[b]** |

| Compound ranking | Compound code | chemical structure | % of SARS-CoV-2 M^pro inhibition (screening at 100 µM) |
|---|---|---|---|
| **49** | **6** | 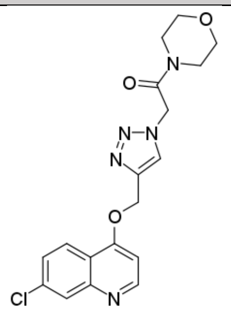 | **10±5** |
| **82** | **7** | 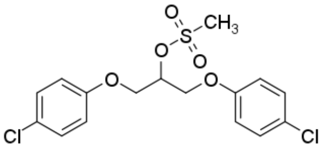 | **23±3[b]** |
| **95** | **8** | 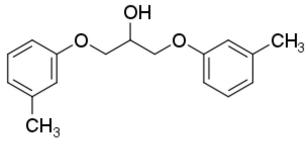 | **19±1** |
| **180** | **9** | 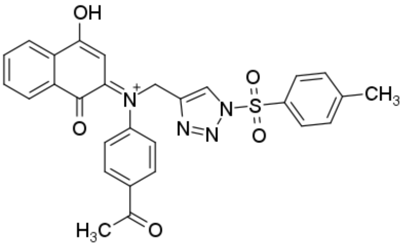 | **50±1** |
| **331** | **10** | 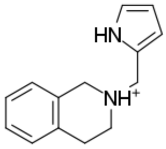 | **0±7** |
| **368** | **11** | 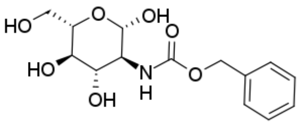 | **11±7** |
| **375** | **12** | | **fluorescent compound[c]** |

| Compound ranking | Compound code | chemical structure | % of SARS-CoV-2 $M^{pro}$ inhibition (screening at 100 μM) |
|---|---|---|---|
| | | 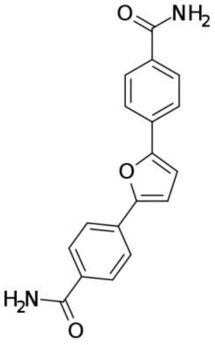 | |
| **378** | **13** | 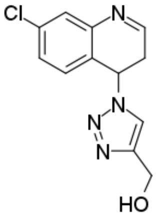 | **27±6** |
| **391** | **14** | 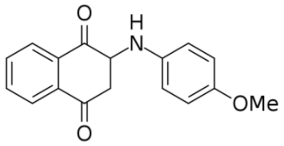 | **20±2[b]** |
| **436** | **15** | 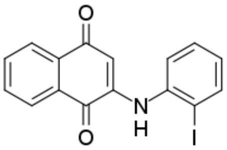 | **59±4[b]** |
| **469** | **16** | 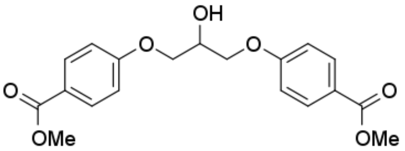 | **50±3** |
| **471** | **17** | | **14±10[b]** |

| Compound ranking | Compound code | chemical structure | % of SARS-CoV-2 $M^{pro}$ inhibition (screening at 100 µM) |
|---|---|---|---|
| | | 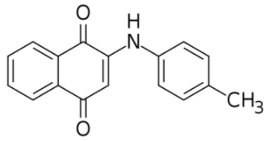 | |
| 476 | 18 | 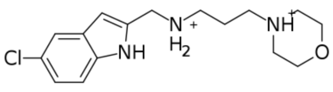 | 0±2 |
| 595 | 19 | 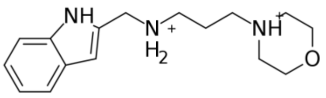 | 0±2 |
| 701 | 20 | 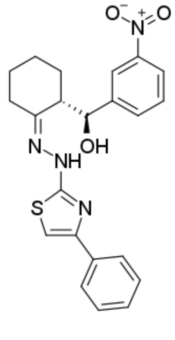 | 16±6[b] |
| 787 | 21 | 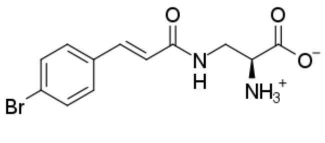 | - |

| Compound ranking | Compound code | chemical structure | % of SARS-CoV-2 $M^{pro}$ inhibition (screening at 100 µM) |
|---|---|---|---|
| **877** | **22** | 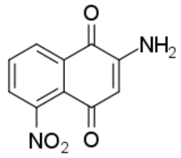 | **1±10** |
| **884** | **23** | 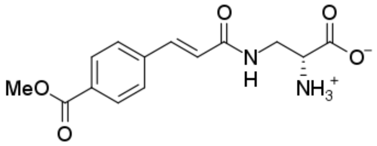 | **-** |
| **892** | **24** | 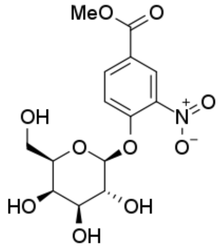 | **0±3** |
| **1053** | **25** | 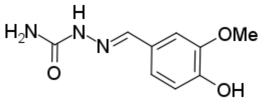 | **57±5** |
| **NA** | **26** | 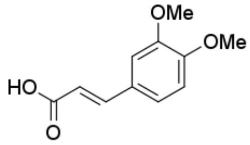 | **fluorescent compound[c]** |
| **NA** | **27** | 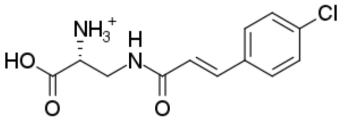 | **0±3** |
| **NA** | **28** | 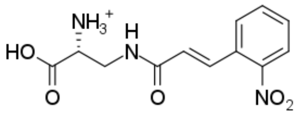 | **0±2** |
| **NA** | **29** | 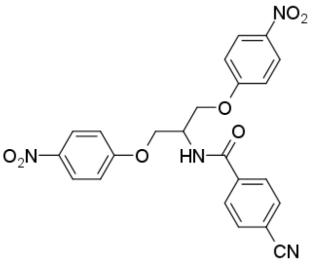 | **22±3[b]** |

| Compound ranking | Compound code | chemical structure | % of SARS-CoV-2 $M^{pro}$ inhibition (screening at 100 µM) |
|---|---|---|---|
| NA | 30 | 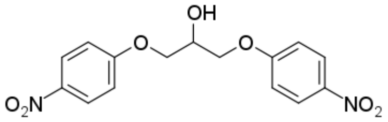 | 31±2[b] |
| NA | 31 | 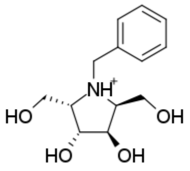 | 0±10 |
| NA | 32 | 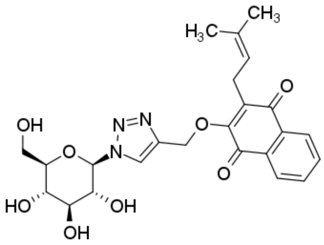 | 50±4 |
| NA | 33 | 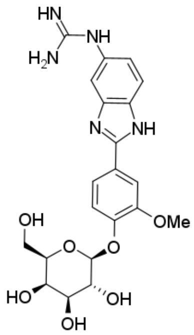 | 0±1[d] |
| NA | 34 | 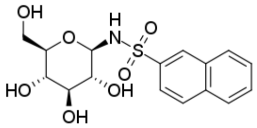 | fluorescent compound[c] |
| NA | 35 | 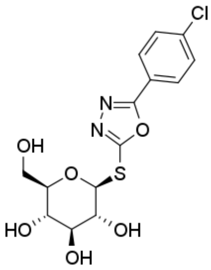 | fluorescent compound[c] |

| Compound ranking | Compound code | chemical structure | % of SARS-CoV-2 $M^{pro}$ inhibition (screening at 100 μM) |
|---|---|---|---|
| **NA** | **36** | 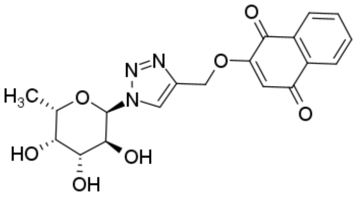 | **fluorescent compound[c]** |
| **NA** | **37** | 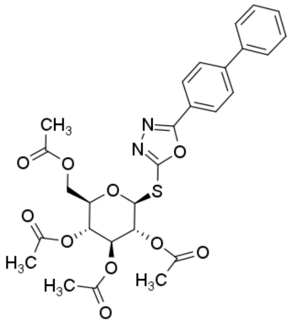 | **0±4** |
| **NA** | **38** | 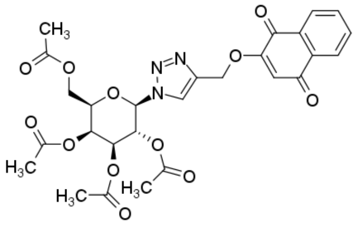 | **fluorescent compound[c]** |
| **NA** | **39** | 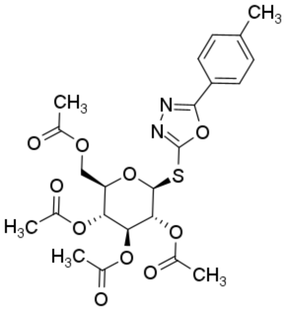 | **20±7** |
| **NA** | **40** | 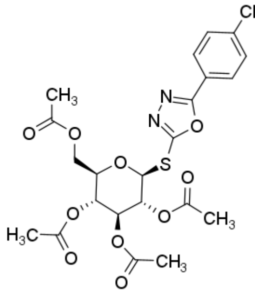 | **0±3** |

| Compound ranking | Compound code | chemical structure | % of SARS-CoV-2 M$^{pro}$ inhibition (screening at 100 µM) |
|---|---|---|---|
| NA | 41 | 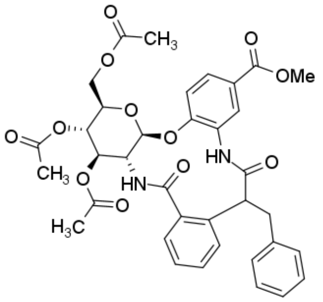 | fluorescent compound[c] |
| Positive control | nirmaltrelvir | 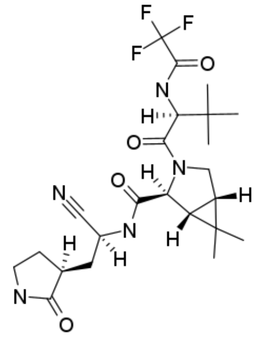 | 96±1 nM[e] |

a Values represent the mean and standard error of the mean (SEM) from two independent experiments, each performed in triplicate. Measurements were conducted in the presence of 100 µM of compound and 0.5% DMSO, unless otherwise specified.

b Compounds tested at 50 µM in the presence of 0.5% DMSO.

c Some compounds exhibited higher intrinsic fluorescence than the fluorogenic substrate, preventing accurate measurement of enzymatic activity.

d Compounds tested at 25 µM in the presence of 0.5% DMSO.

*NA These compounds were not included in the virtual screening and correspond to analogs of hits from the BraCoLi library.

e Compounds tested at 100 nM in the presence of 0.5% DMSO.

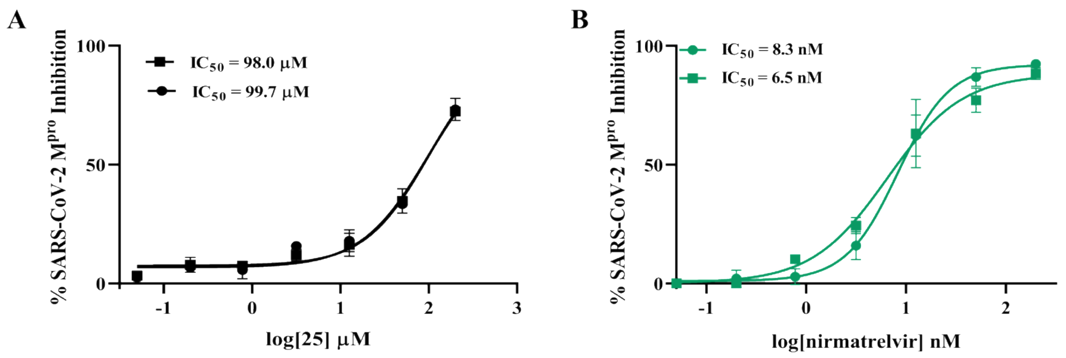


**Figure S2.** Determination of $IC_{50}$ for SARS-CoV-2 $M^{pro}$ inhibition by compound **25** and nirmatrelvir. Concentration-response curves for SARS-CoV-2 $M^{pro}$ inhibition by (A) **25** (concentration range: 0.15–200 μM) and (B) nirmatrelvir (concentration range: 0.05-200 nM). Each $IC_{50}$ curve represents an independent experiment performed in triplicate following preincubation with the enzyme (40 nM), followed by addition of the fluorogenic substrate MCA-AVLQSGFR-Lys(Dnp)-Lys-NH2 (10 μM).

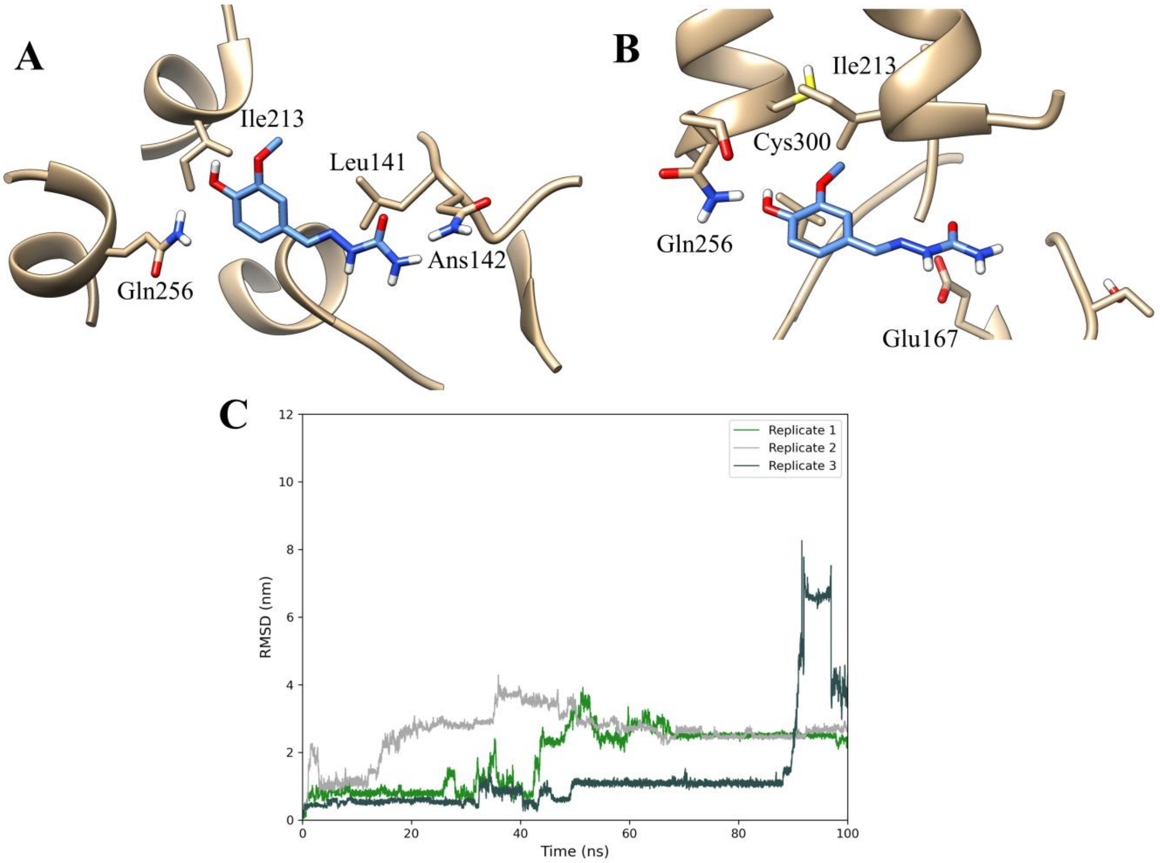


**Figure S3**. Predicted binding mode of compound **25** at the allosteric site of SARS-CoV-2 M$^{pro}$ using DockThor VS mode (A) and Standard mode (B). Both poses were used as the initial frame for the MD simulations. (**C**) Root mean square deviation RMSD (nm) of the best-ranked binding pose of compound **25**, obtained from the DockThor standard mode, over a 100 ns molecular dynamics simulation per replicate.
