## Supplementary material for "Discovery of Semicarbazone and Thiosemicarbazone Analogs as Competitive SARS-CoV-2 Virus Main Protease (M^pro^) Inhibitors": Spectral data

### Compound 1

#### Infrared spectrum (ATR)

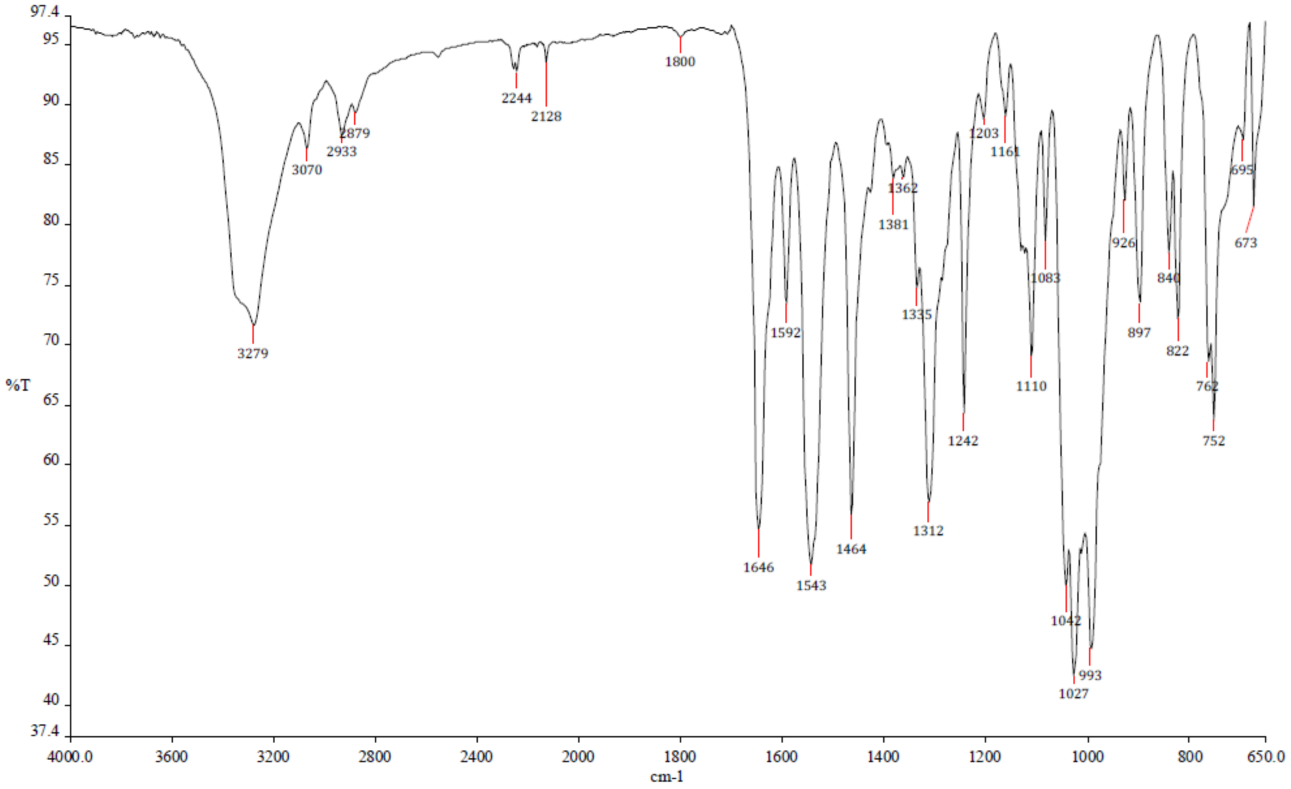


#### [1]H NMR spectrum (400 MHz, DMSO-$d_6$)

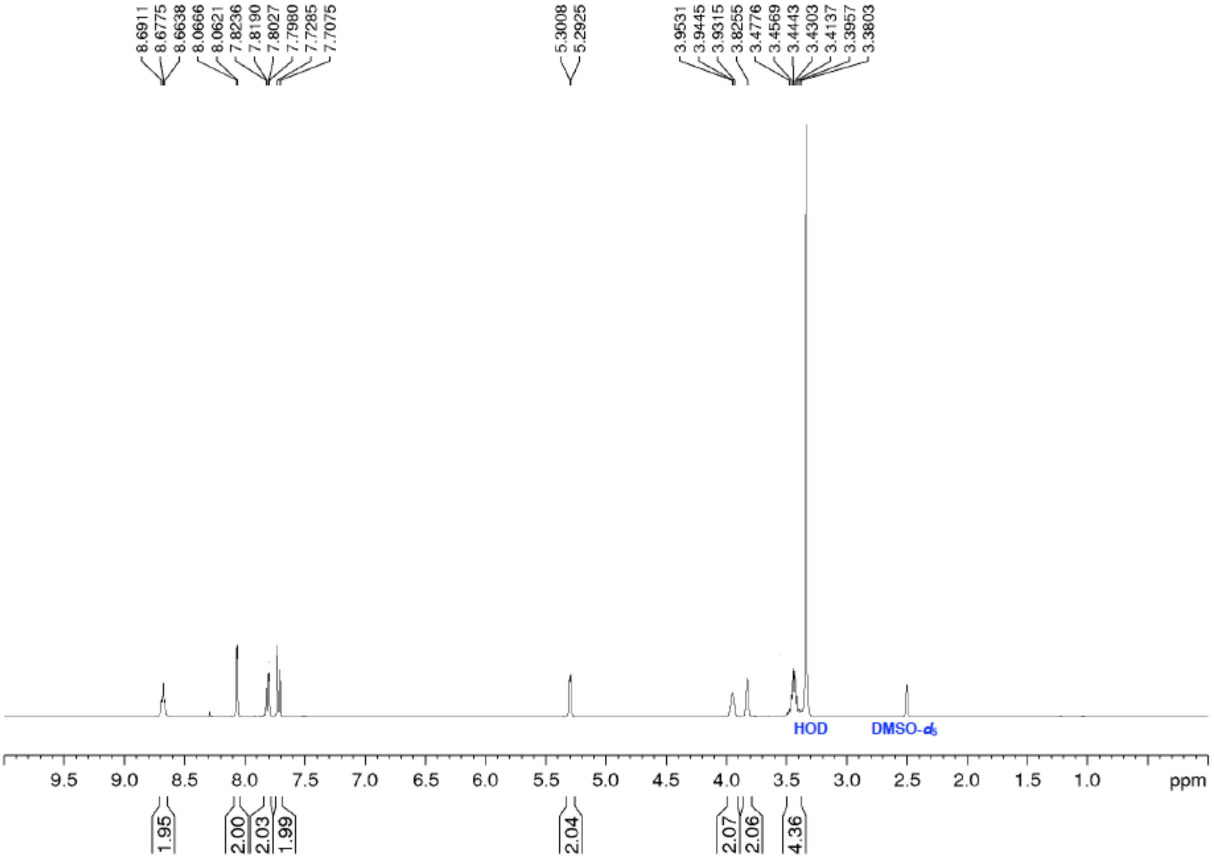

**$^{13}$C NMR spectrum (100 MHz, DMSO-$d_6$)**

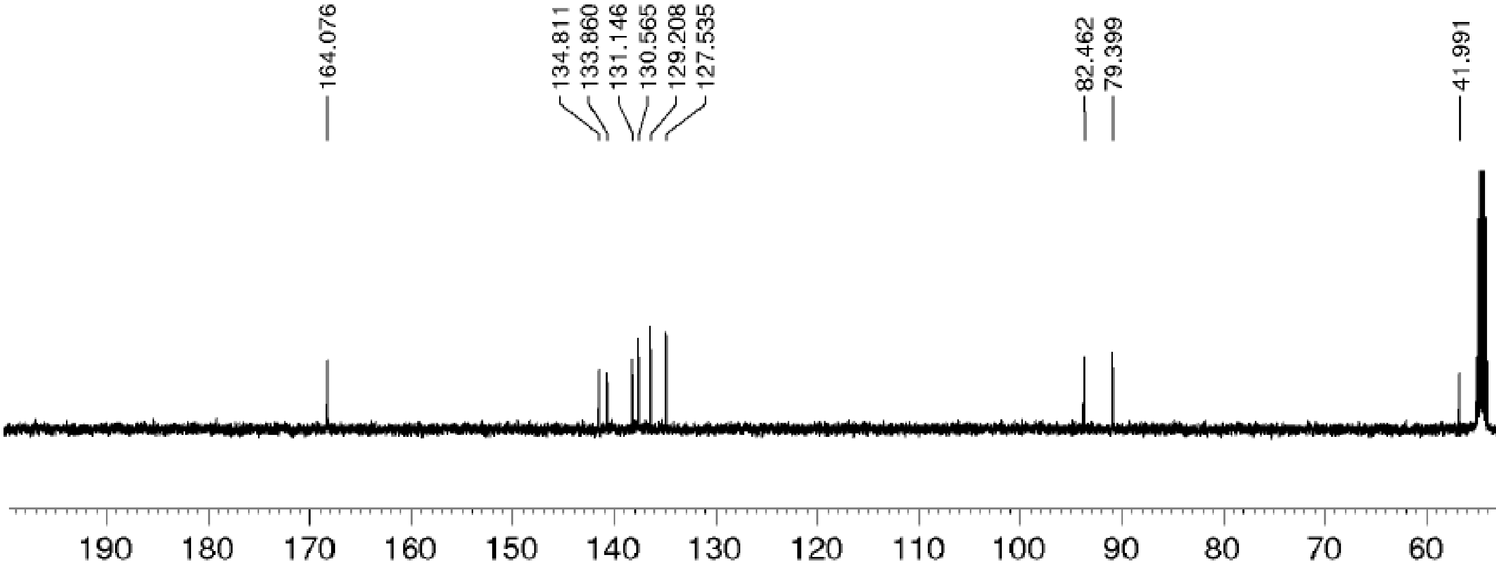

### Compound 5

#### $^{1}$H NMR spectrum (200 MHz, DMSO-$d_6$)

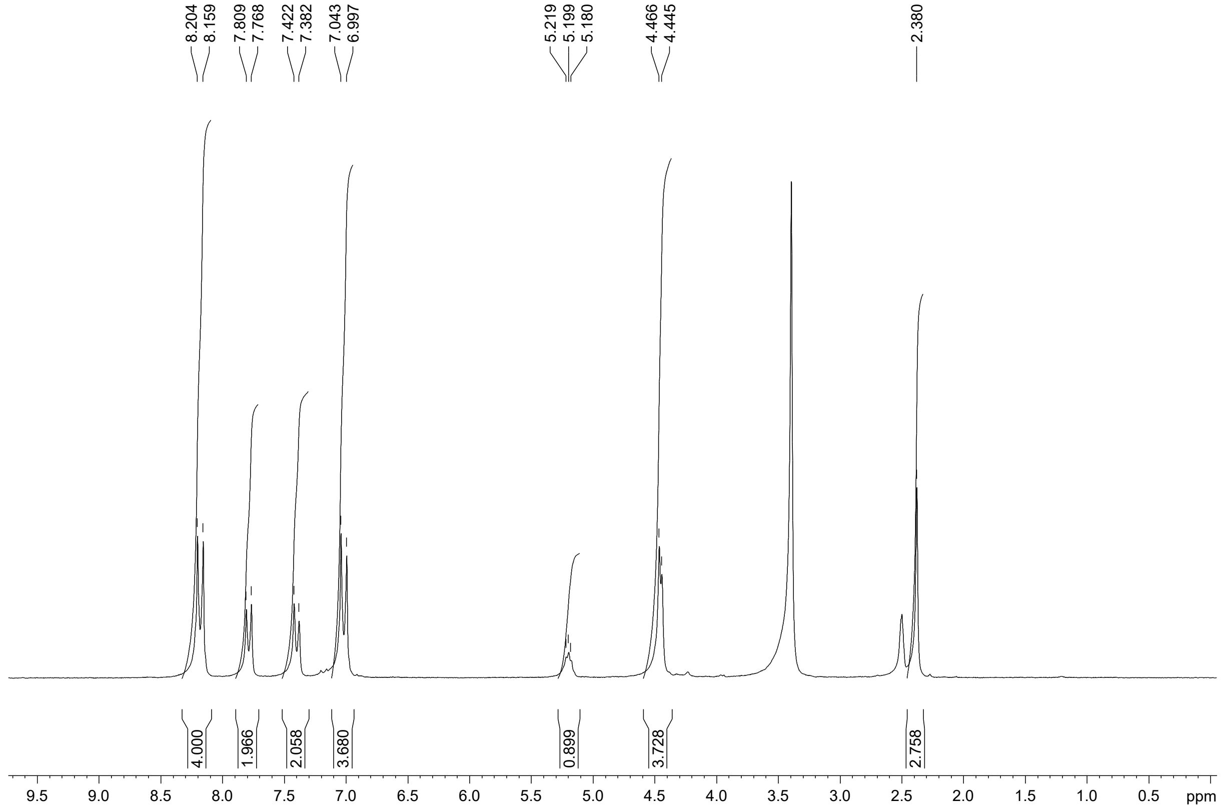


#### $^{13}$C NMR spectrum (50 MHz, DMSO-$d_6$)

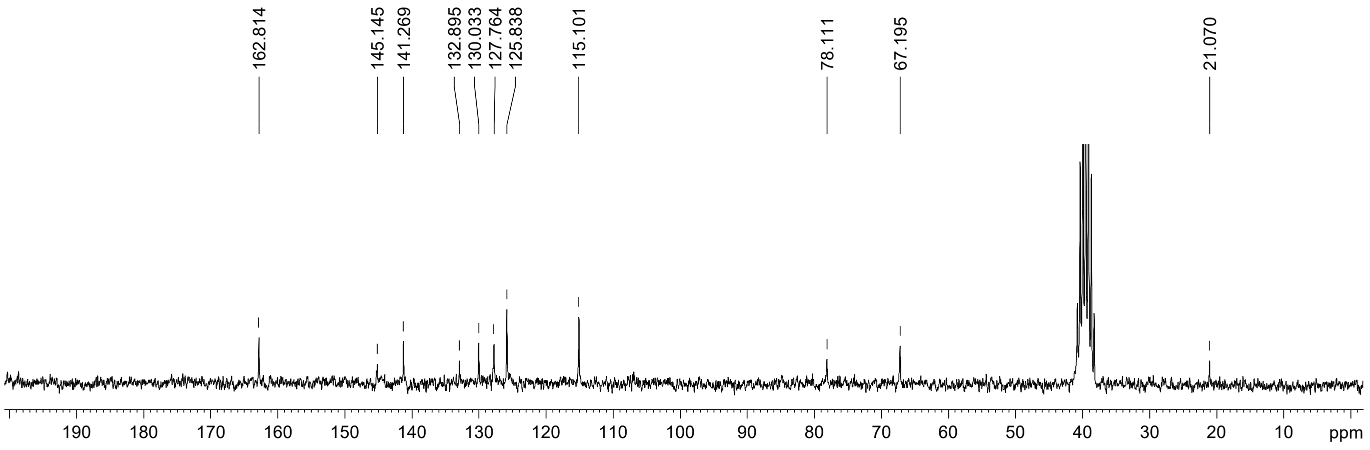

### Compound 21

#### [1]H NMR spectrum (400 MHz, $D_2O$/TFA)

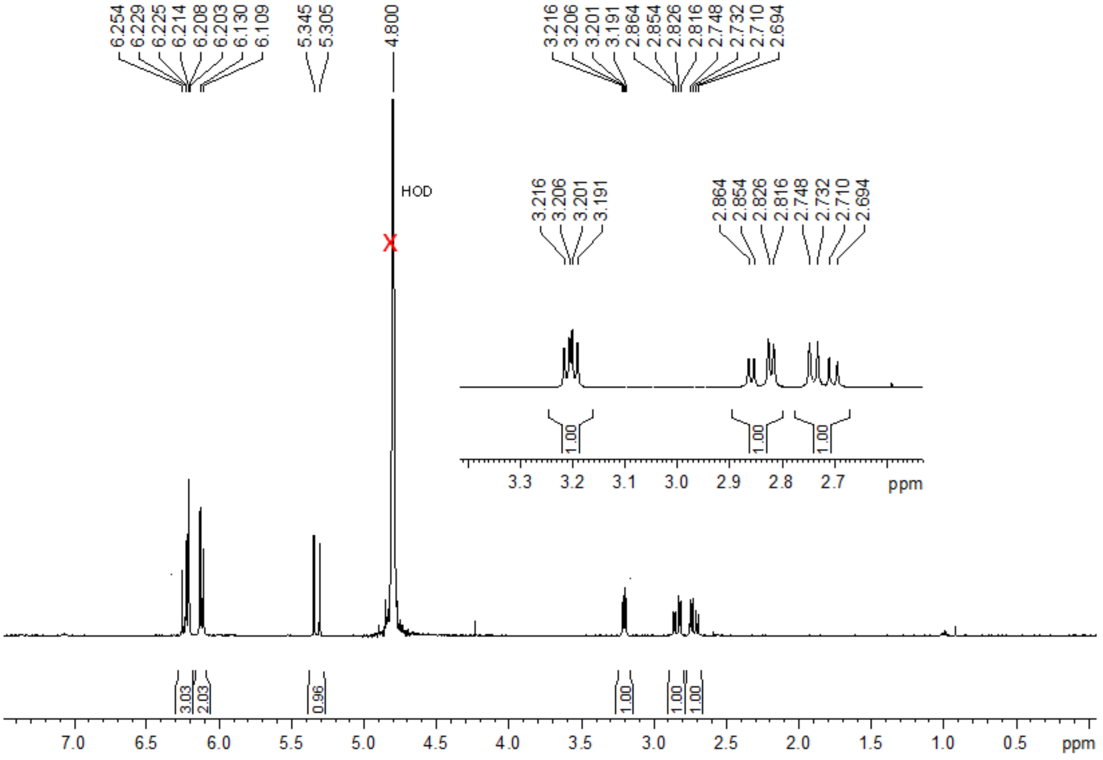


#### [13]C NMR spectrum (100 MHz, $D_2O$/TFA)

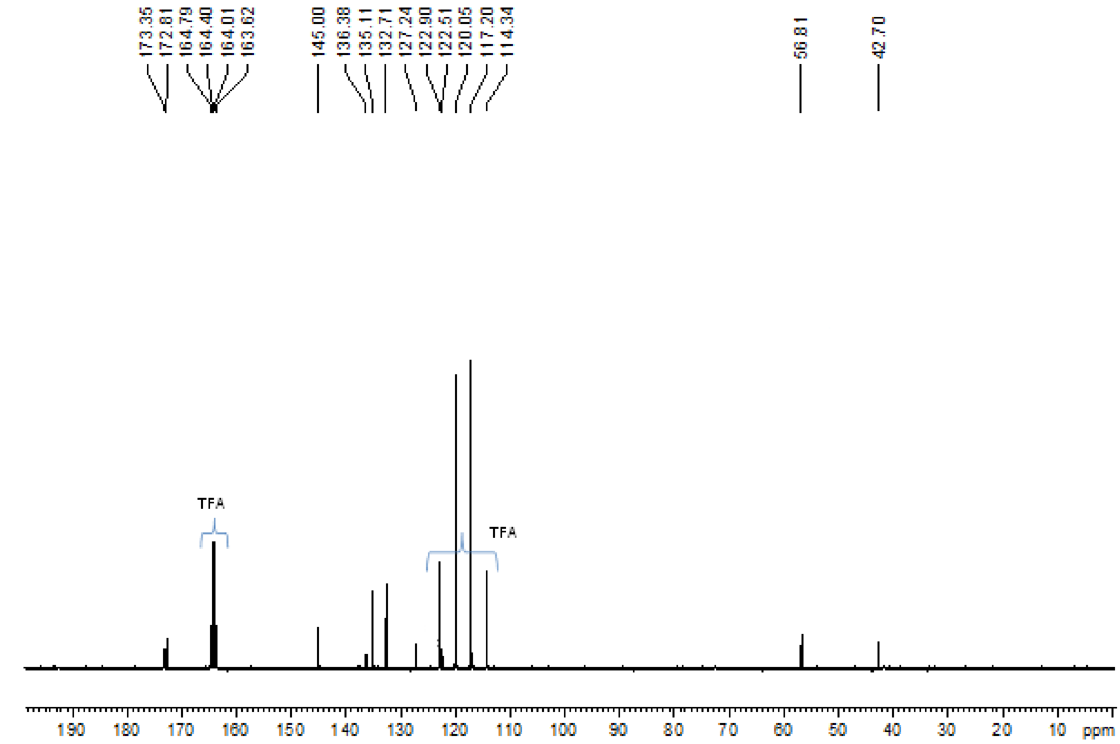


#### Mass spectrum (ESI+)

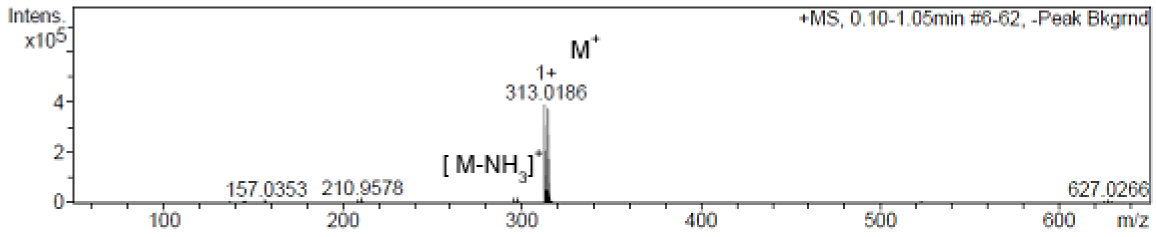

**Compound 23**

**$^{1}$H NMR spectrum (400 MHz, $D_2O$/TFA)**

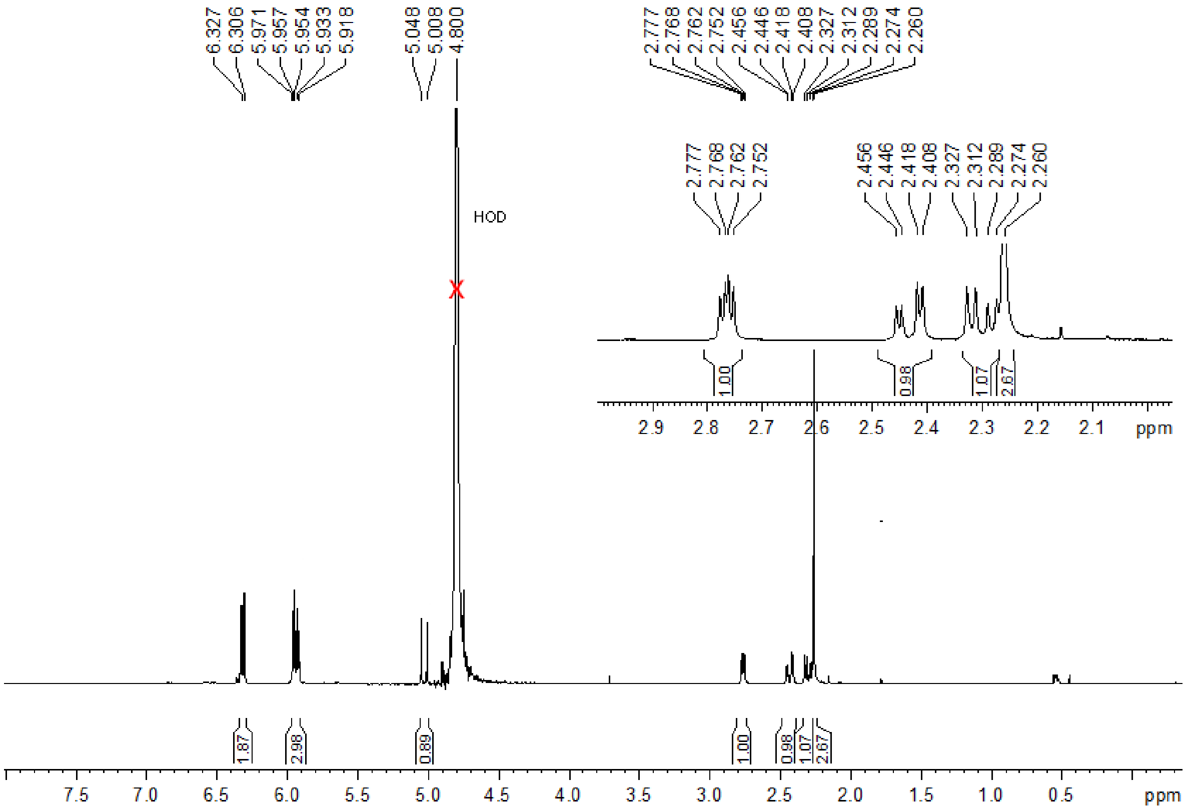


**$^{13}$C NMR spectrum (100 MHz, $D_2O$/TFA)**

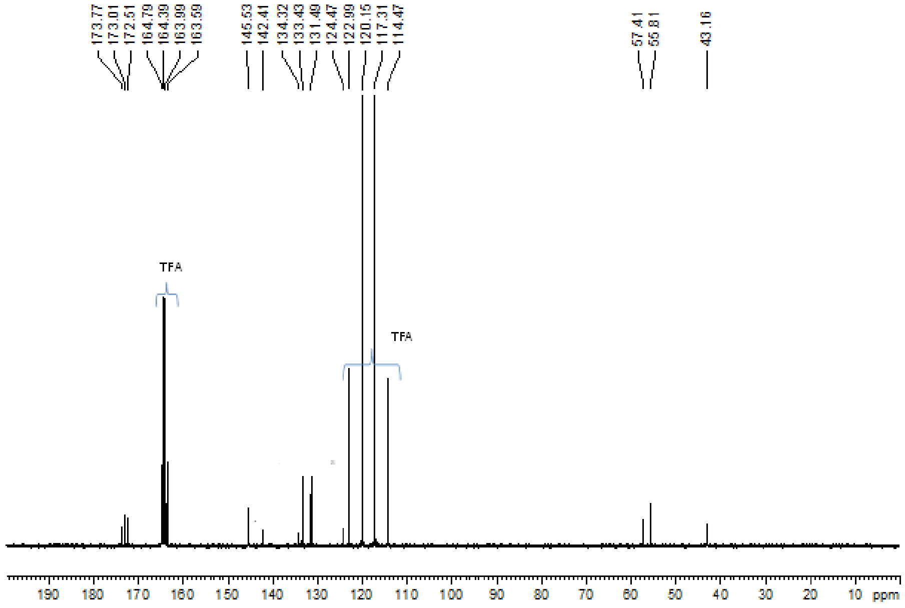


**Mass spectrum (ESI+)**

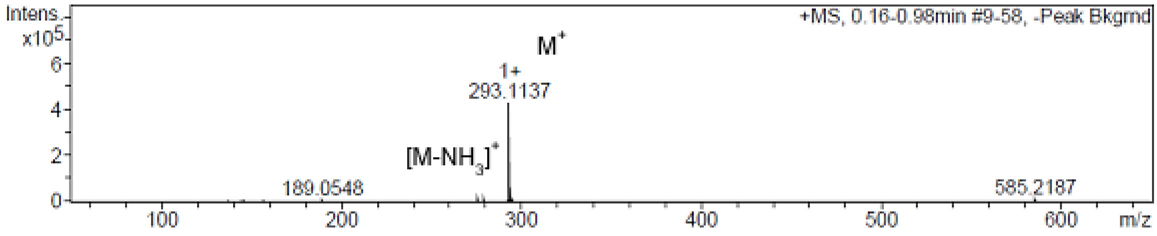

### Compound 27

#### $^{1}H$ NMR spectrum (400 MHz, $D_2O$/TFA)

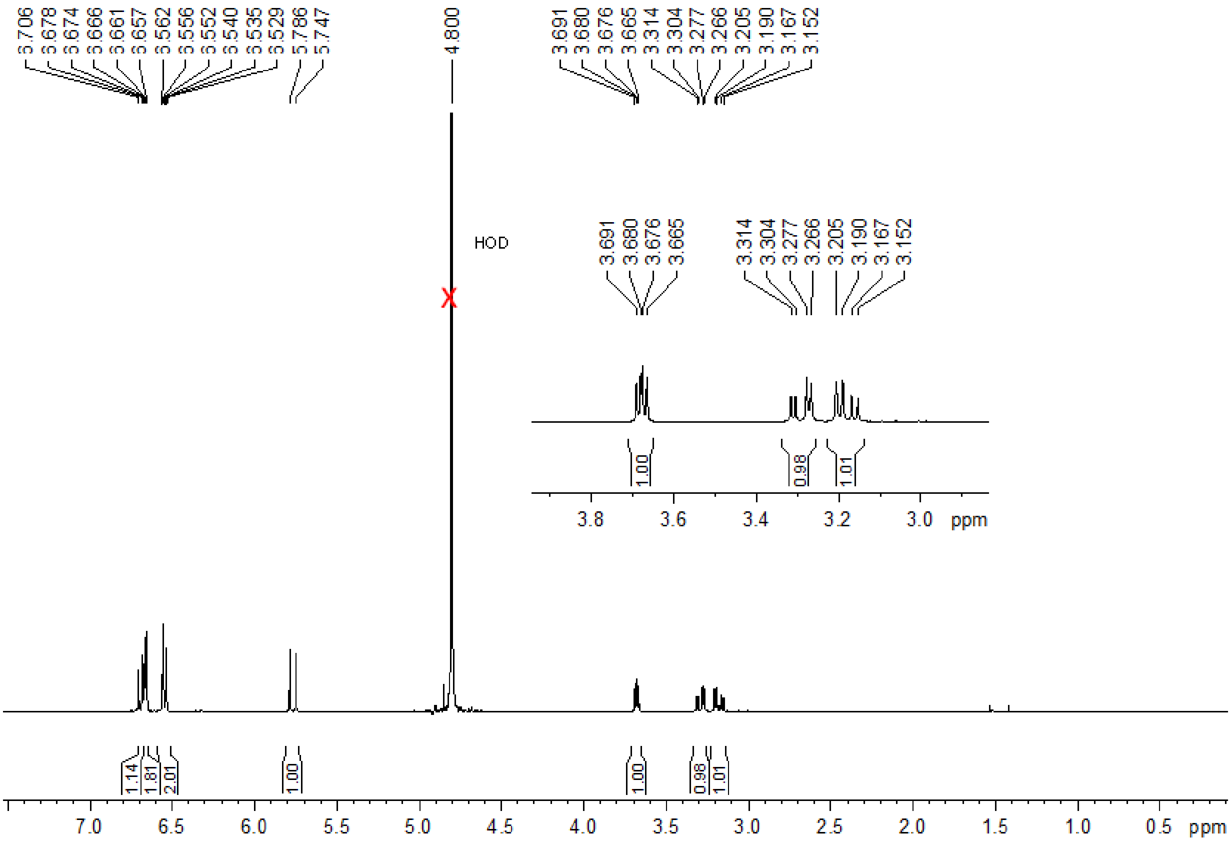


#### $^{13}C$ NMR spectrum (100 MHz, $D_2O$/TFA)

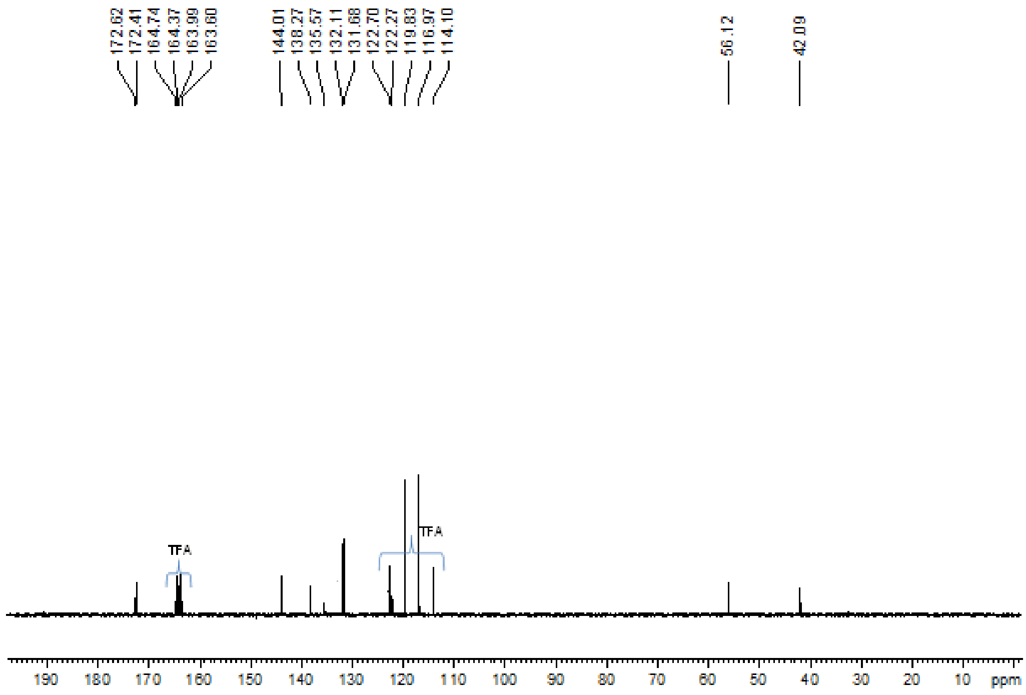


#### Mass spectrum (ESI+)

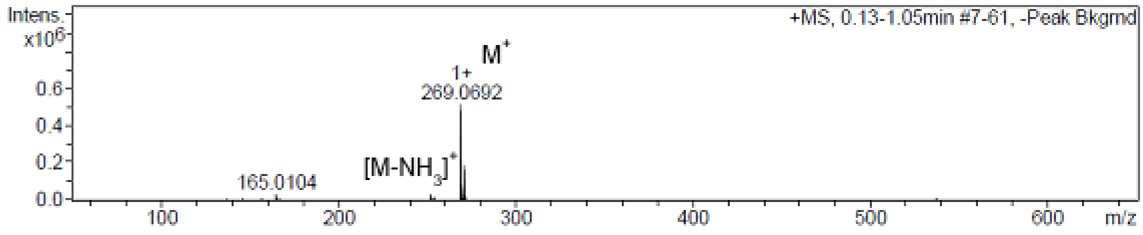

### Compound 28

#### $^{1}H$ NMR spectrum (400 MHz, $D_2O$/TFA)

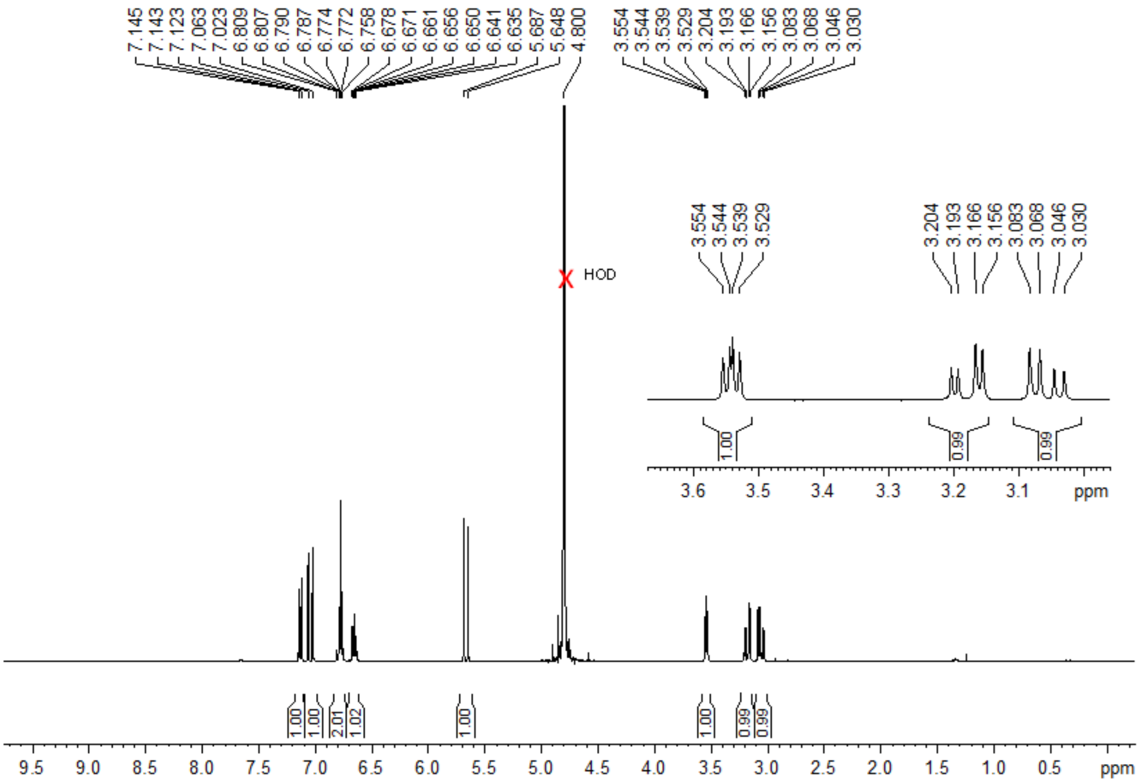


#### $^{13}C$ NMR spectrum (100 MHz, $D_2O$/TFA)

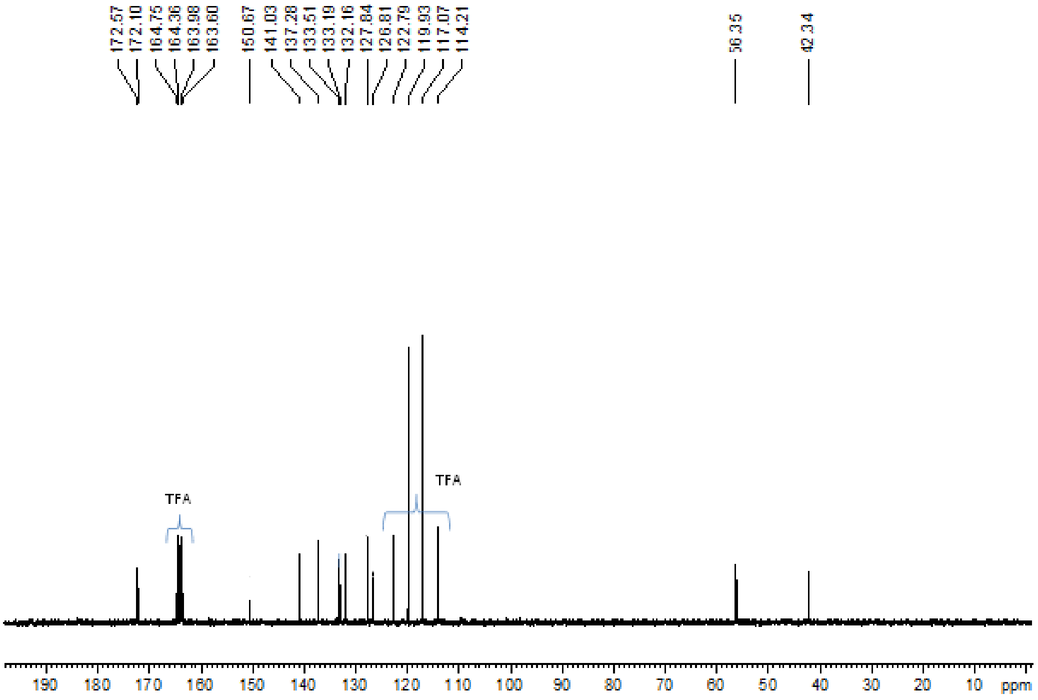


#### Mass spectrum (ESI+)

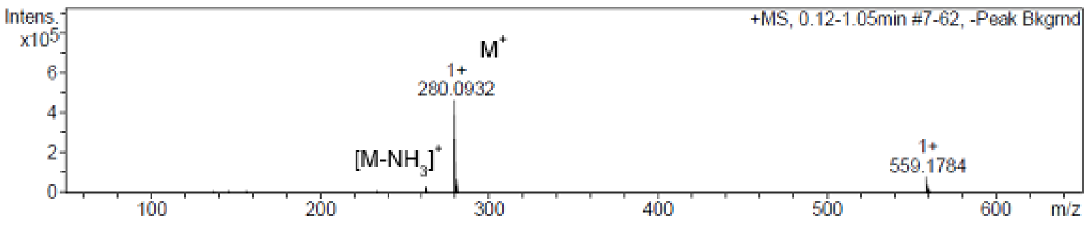

**Compound 32**

**Infrared spectrum**

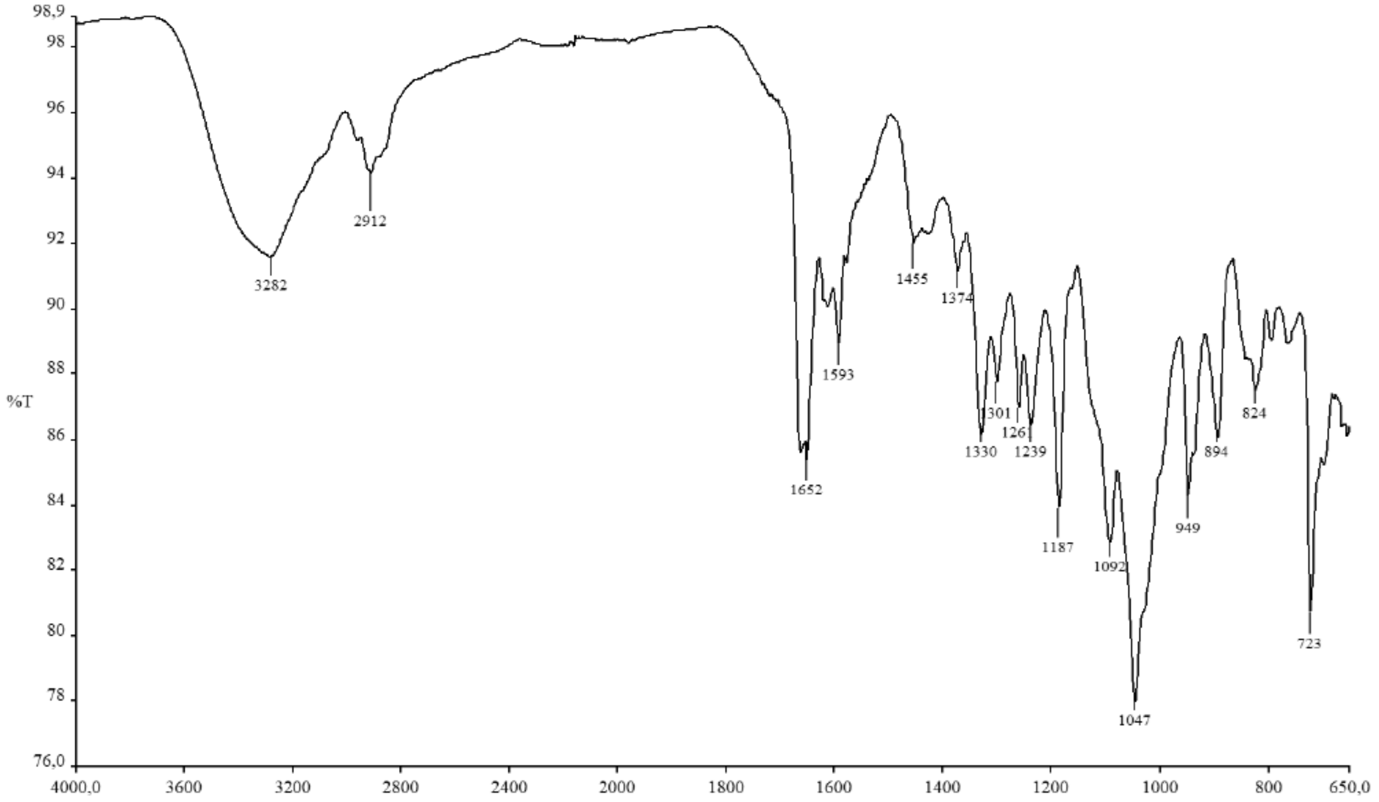

**[1]H NMR spectrum (400 MHz, DMSO-$d_6$)**

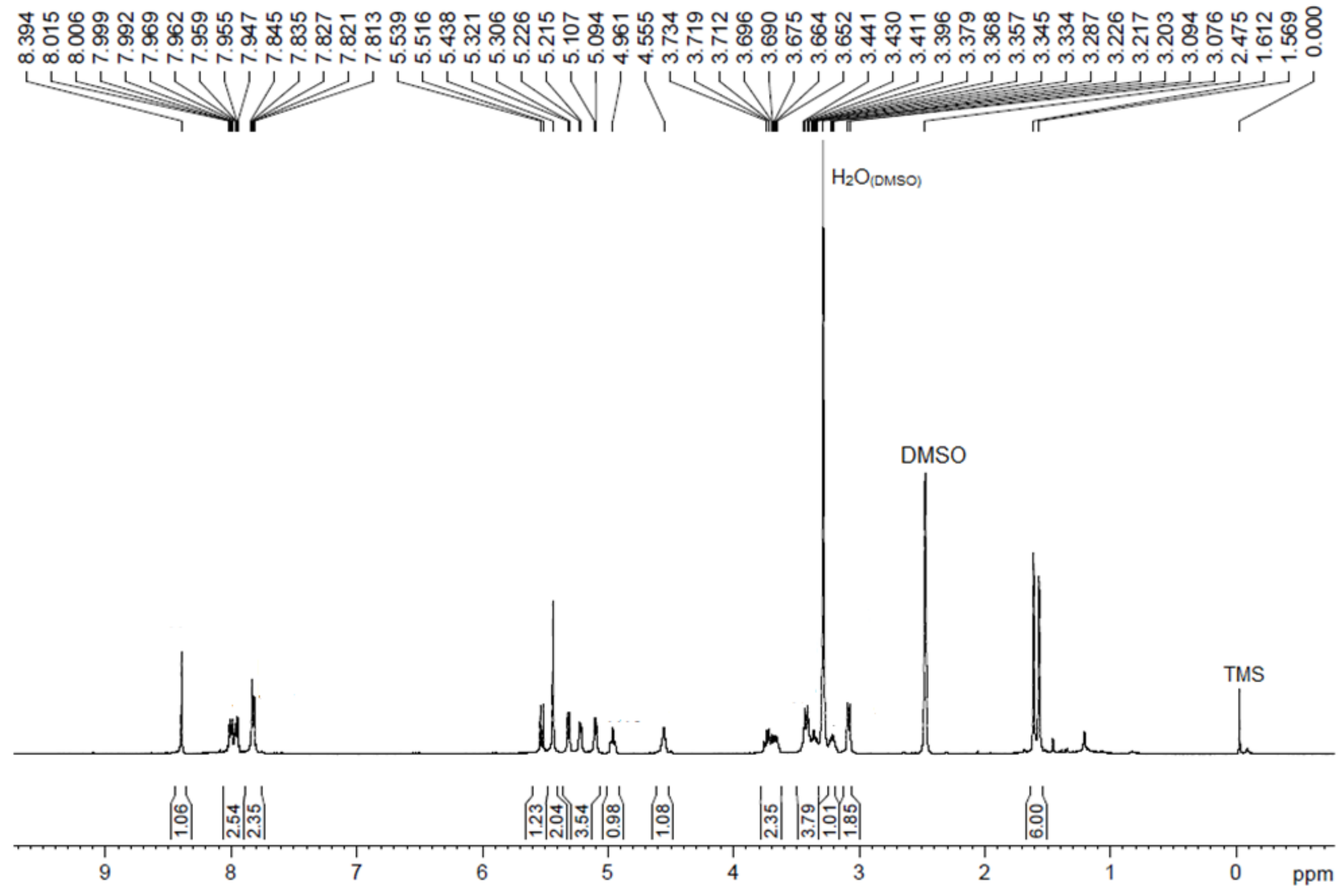

### $^{13}C$ NMR and DEPT135 spectra (100 MHz, DMSO-$d_6$)

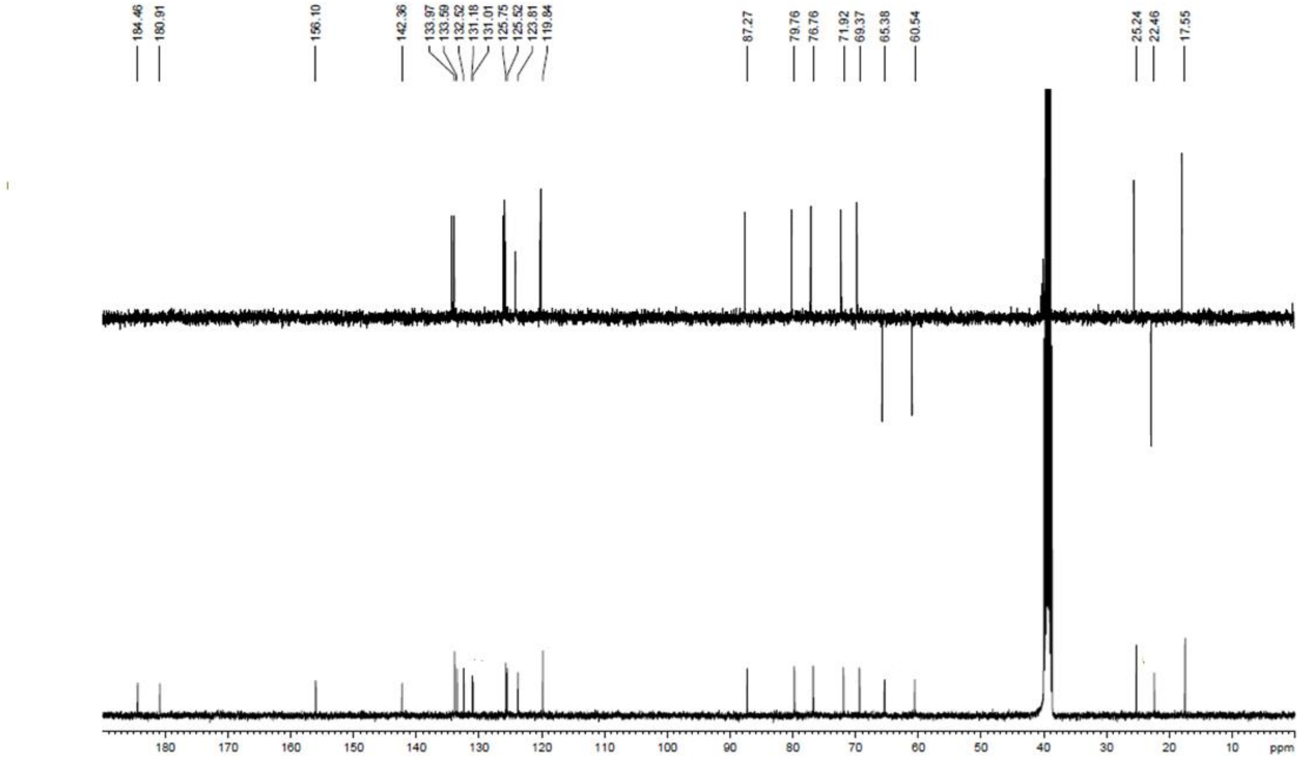


### UPLC analysis

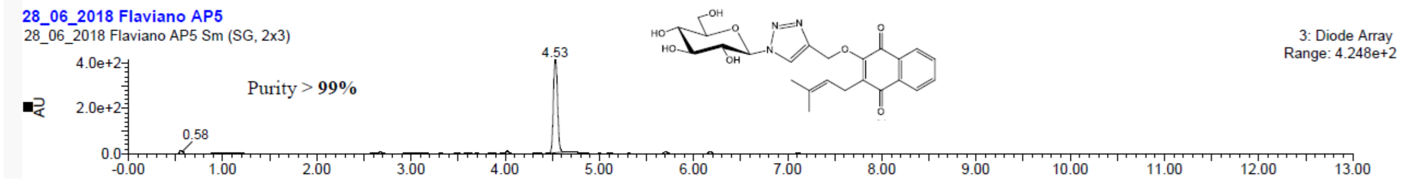


### Mass spectrum (ESI+)

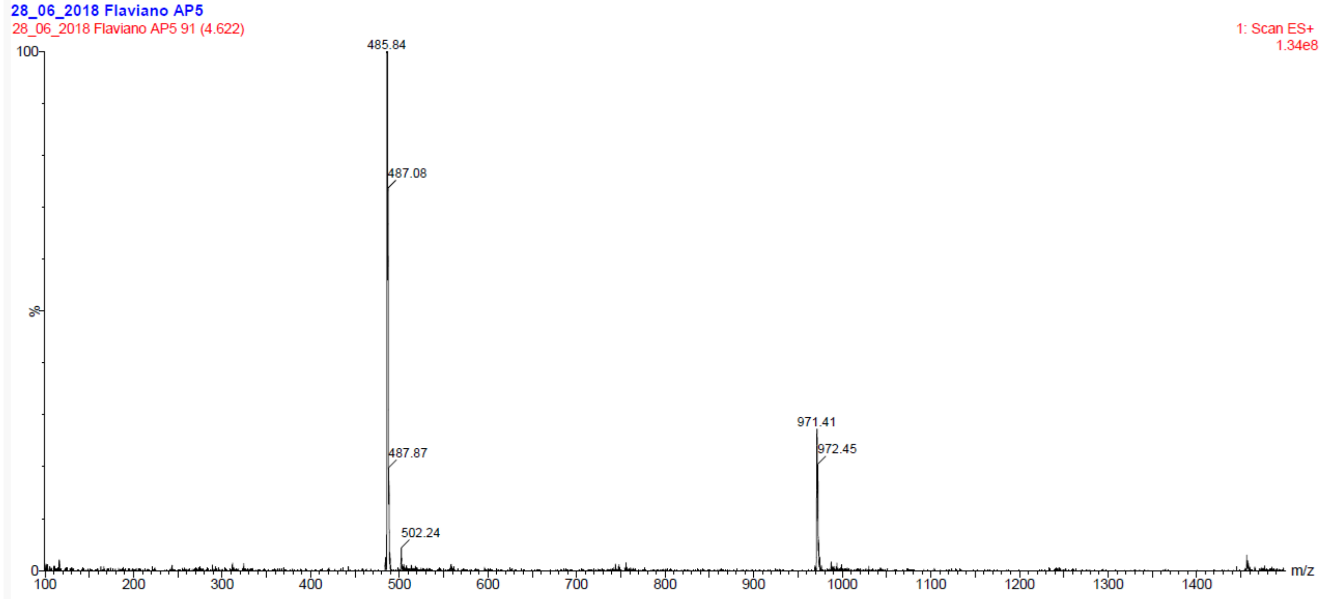

**Compound 41**

**$^{1}$H NMR spectrum (400 MHz, DMSO-$d_6$)**

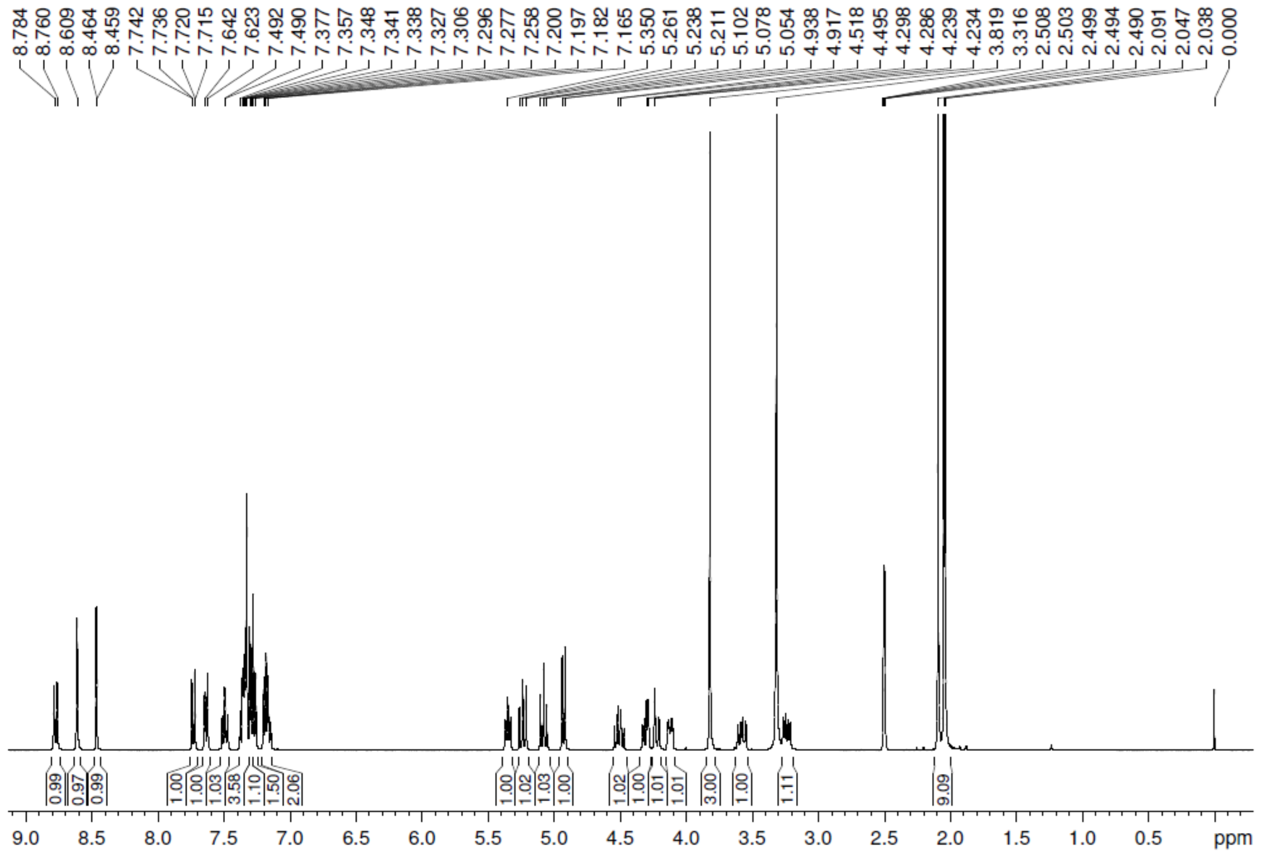

**$^{1}$H NMR spectrum (400 MHz, DMSO-$d_6$) - expansion**

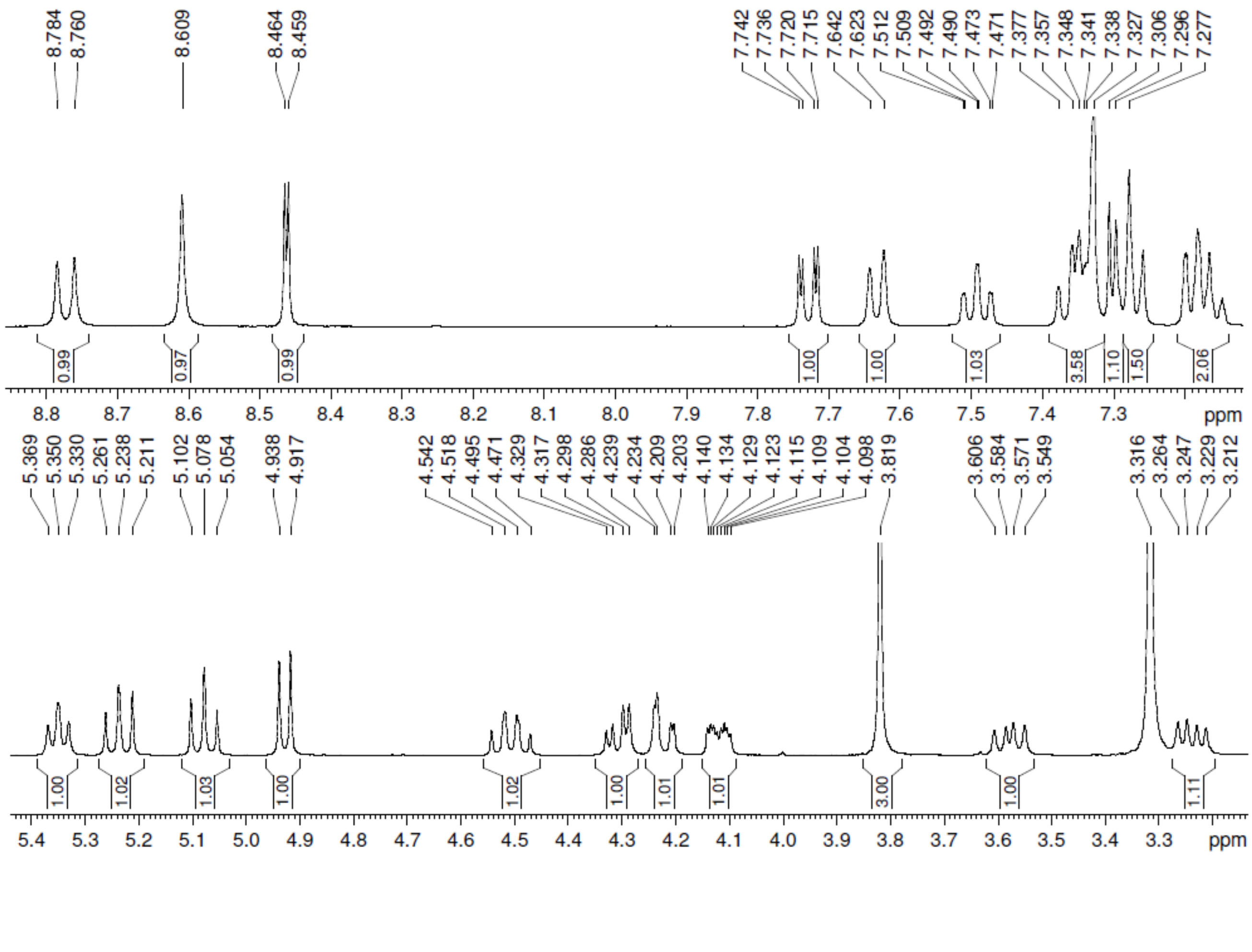

### $^{13}C$ NMR spectrum (100 MHz, DMSO-$d_6$) and expansion

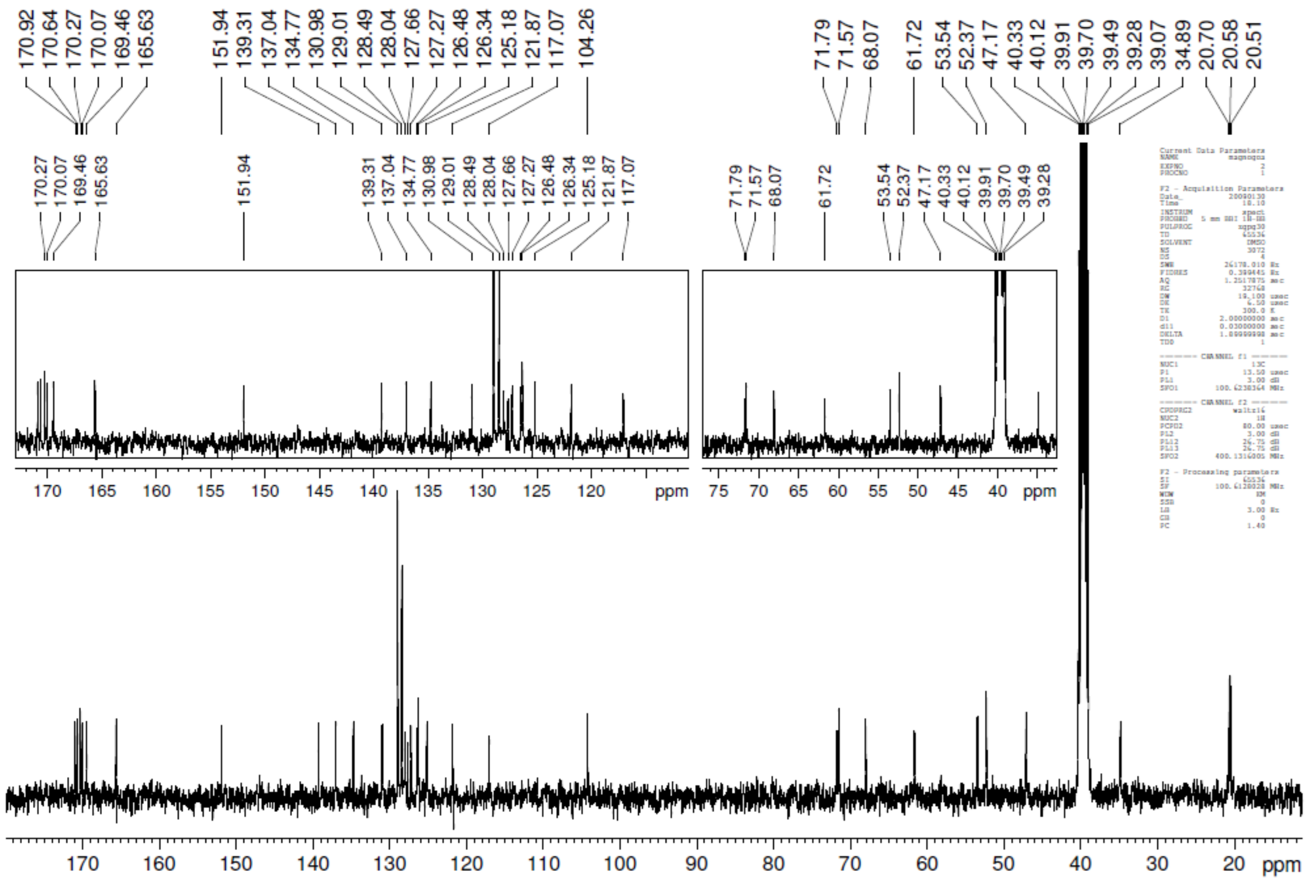


### Mass spectrum (TOF MS ES+)

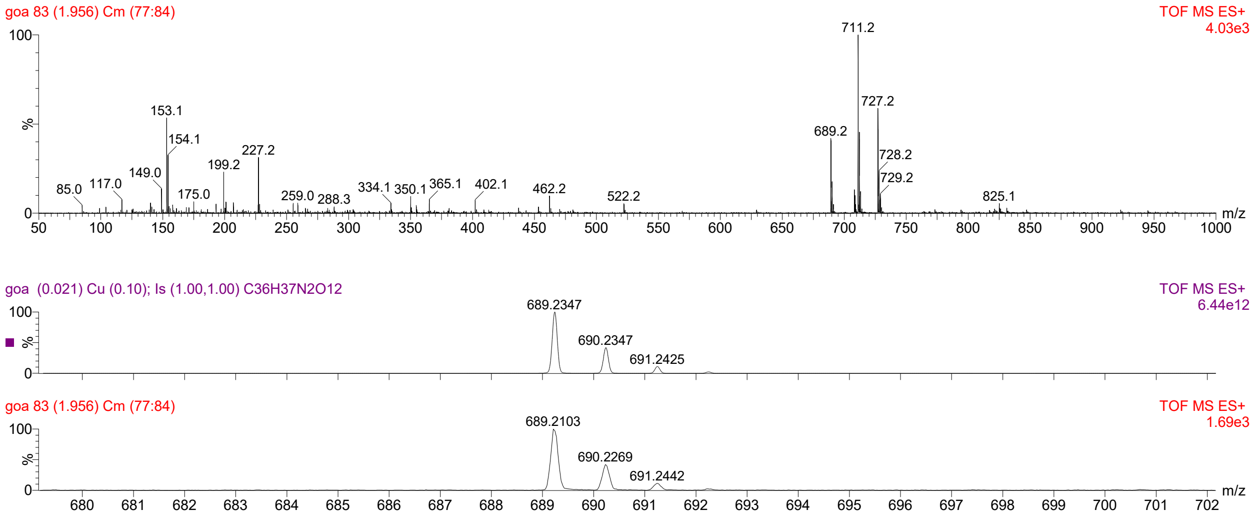
